# Pharmacological inhibition of an epigenetic regulator during the formation of extinction memory reveals sensory cortical and subcortical codes for the specificity of salient cues

**DOI:** 10.1101/2020.07.31.230524

**Authors:** Elena K. Rotondo, Kasia M. Bieszczad

## Abstract

Using a sound-reward extinction paradigm in male rats, we reveal both cortical and subcortical sensory codes for the cue-specificity of memory. In the auditory cortex, re-tuning narrowed frequency receptive field bandwidth, yielding more precise extinction behavior cued by acoustic frequency. Subcortical signals revealed in the auditory brainstem response (ABR) showed decreases in amplitude of select components of the ABR wave. Interestingly, treatment with an inhibitor of histone deacetylase 3 (HDAC3-i) facilitated both auditory cortical tuning bandwidth changes and ABR changes that were frequency-specific to the extinguished signal sound. Moreover, both changes were correlated to each other and with highly precise extinction memory at the level of behavior. Thus, we show for the first time that HDAC3 regulates the specificity of sensory features consolidated in extinction memory. Overall, the dynamics of auditory system plasticity associated with sound-specific extinction are complex. Changes in ABR amplitude induced by sound-reward learning disappeared after extinction, while changes in ABR slope that were initially induced by sound-reward learning were maintained through extinction. Moreover, plasticity of cortical re-tuning emerged only after extinction learning. HDAC3-i applied after extinction training sessions enabled sensory system plasticity to encode the extinguished sound with higher acoustic specificity (compared to vehicle controls). Both cortical and subcortical response changes to sound became unusually “tuned-in” to the acoustic frequency that had been presented under extinction conditions. Thus, HDAC3 regulates how specifically sensory features of experience are encoded into long-term memory and may exert its behavioral effects via multiple coding strategies along sensory system pathways.

**SIGNIFICANCE STATEMENT:** Epigenetic mechanisms have recently been implicated in memory and information processing. Here, we use a pharmacological inhibitor of histone deacetylase 3 (HDAC3) in a sensory model of learning to reveal, for the first time, its ability to enable unusually precise extinction memory. In so doing, we uncover neural coding strategies for memory’s “specificity” for sensory cues. Extinction induced multiple forms of change at different levels of sensory processing, which highlights the complexity of extinction memory encoding. HDAC3 appears to coordinate effects across sensory levels that determine specific cue saliency for behavior. Thus, epigenetic players may gate how sensory information is stored in long-term memory and their manipulation can be leveraged to reveal neural coding mechanisms for sensory detail in memory.

## 1. INTRODUCTION

Epigenetic mechanisms are now known to be ubiquitous in long-term memory processes. Previous investigations have shown one variety of epigenetic regulation by class I histone deacetylases (HDACs)—including the most prominent HDAC expressed in the adult brain, HDAC3—can enable long-term memory (LTM) formation in a variety of behavioral tasks and species (Bredy et al., 2007; Lattal et al., 2007; Stefanko et al., 2009; Malvaez et al., 2010; Wang et al., 2010; Malvaez et al., 2013; Raybuck et al., 2013; Bowers et al., 2015; Hitchcock et al., 2019; Phan et al., 2017) including humans (Gervain et al., 2013). However, recent work using a sensory model of learning, memory and experience-dependent sensory neuroplasticity has shown a much more significant role of HDAC3 on LTM beyond facilitating its formation *per se*. Treatment with a selective HDAC3-inhibitor during auditory associative learning appears to induce a dramatic change in information processing to alter what and how much acoustic information becomes encoded to produce persistent and vivid memory in the long-term (Bieszczad 2015, Shang 2019, Rotondo 2020). In the auditory system, this is realized by a reorganization of sound-evoked responses to increase the neural responsivity and selectivity to the specific training sounds that are later remembered with high sensory precision and accuracy, which we operationally define as “specificity” (Bieszczad 2015, Shang 2019, Rotondo 2020). The present study used an established auditory model of memory formation with an HDAC3-inhibitor to determine whether the impact to gate the formation of memory with sensory specificity is a general function of HDACs. To do so, we asked whether HDAC3-inhibition during *extinction* of a sound-reward association would promote acoustic specificity in subsequent extinction memory. This is the first study to examine cued memory specificity in extinction. Extinction memory provides a complementary contrast to initial sound-cued reward memory in which the associative link to reward and the learned behavioral output is in direct opposition to that of prior learning. However, an extinguished sound still has a salient meaning, which permits the study of common mechanisms for memory specificity *per se*. Addressing this question will determine whether HDAC3 functions to enhance the sensory specificity of memory regardless of associative value of the sensory cue (i.e., whether it signals reward or no-reward) and the associated behavioral response (i.e., activating or inhibiting a behavioral action). Further, this strategy provides a unique opportunity to link particular forms of auditory system reorganization with the key characteristics of memory, such as its persistence in time, its associative strength, or its specificity (Weinberger, 2015). If the auditory system reorganizes in same way in extinction as it does for sound-reward acquisition to increase the acoustic specificity of subsequent behavior, then that coding principle is not related to associative value, nor the direction of behavior, but to memory’s acoustic specificity *per se*. Since sensory system contributions to memory include subcortical substrates of information encoding with likely cortical-subcortical interactions (i.e., as in Rotondo & Bieszczad 2020), we probe the neural sequelae of HDAC3-inhibition on auditory memory in both the auditory cortex (ACx) and the auditory brainstem response (ABR). As such, we show for the first time that the use of HDAC manipulations are prime tools to discover the neural coding strategies for sensory information storage in sensory systems, while also uncovering a key role that HDACs play in forming the characteristics of longterm memory.

## 2. Methods

### 2.1 Subjects

A total of 20 adult male Sprague-Dawley rats (275-300 g on arrival; Charles River Laboratories, Wilmington MA) were used (RGFP966: n=7; vehicle: n=7; no extinction control: n=6) in behavioral and electrophysiological procedures for the main set of extinction experiments. In sum, these rats represent 3 separate groups: (1) Vehicle-treated: rats that received vehicle injections during extinction training, (2) RGFP966-treated: rats that received RGFP966 injections during extinction training, and (3) No-Extinction Control: rats that received tone-reward training, but did not undergo extinction and did not receive injections. This last group of rats were used for baseline (pre-extinction) comparison of cortical electrophysiology. An additional 13 adult male Sprague-Dawley rats (275-300 g on arrival; Charles River Laboratories, Wilmington MA) were used for a control experiment in a task that targeted initial sound-reward learning: (RGFP966: n = 6; vehicle: n = 7). Separate behavioral and neural datasets from these animals have been published (Rotondo & Bieszczad, 2020); novel neural analyses are presented here. All animals were individually housed in a colony room with a 12-hour light/dark cycle. Throughout behavioral procedures, rats were water-restricted, with daily supplements provided to maintain at ~85% free-drinking weight. All procedures were approved and conducted in accordance with guidelines by the Institutional Animal Care and Use Committee at Rutgers, The State University of New Jersey.

### 2.2 Behavioral Approach

All behavioral sessions were conducted in instrumental conditioning chambers within a sound-attenuated box. All subjects initially learned how to press a lever for water reward in five ~45-minute barpress shaping sessions. This phase of training assured that all animals could acquire the procedural aspects of the task (i.e., bar-pressing for rewards) before any sounds were introduced.

Next, rats underwent ***tone-reward training*** in a ***multiple*** tone detection task, in which they learned to associate five different pure tone signals (2.17, 4.2, 5.0, 5.946, and 11.5 kHz, 8s, 60 dB) with the operant reward. The purpose of this task was to create a frequency-general behavioral gradient to set the stage for detecting frequency-specific extinction. Responses in the presence of the signal tone were rewarded, while responses during the variable intertrial interval (ITI; mean = 15 s, range = 5-25 s) triggered a visual error signal and a 6 s time-out that extended the time until the next tone trial. All rats were trained to performance criteria, where on average 70% of bar presses occurred in the presence of the signal tone for 2 consecutive days.

Forty-eight hours following the final tone-reward training session, rats underwent the first ***single-tone extinction training*** session. Initially, there were 10 rewarded trials (2 of each trained tone frequency) separated by a variable ITI (mean = 15 s, range = 5-25 s) to ensure rats pressed to all tones equally. Subsequently, there were 25 unrewarded extinction trials in which only the 5.0 kHz signal tone was presented, separated by a variable ITI (mean = 20 s, range = 10-30 s). No responses were reinforced. Forty-eight hours following the first extinction training session, rats underwent a second extinction training session in which there were 15 unrewarded extinction trials with the 5.0 kHz signal tone. Because the task does not explicitly require discrimination among tones, it allows for individual differences in the strategies used to learn. Thus, rats could learn and remember the actual extinction training sound frequency and inhibit responding selectively, or they could learn and remember to inhibit responding to any sound *per se*.

Forty-eight hours following the final extinction training session, rats were tested in an ***Extinction Memory Test*** to reveal the strategy they had used to learn and remember the task: frequency-specific or frequency-general? The Extinction Memory Test was used to determine the degree of memory specificity for the extinguished signal tone frequency. In the Extinction Memory Test, rats are presented with the 5.0 kHz extinguished signal tone as well as the 4 non-extinguished signal tones, representing tones that are “nearby” neighbors (+/− 0.25 octaves) to the extinguished 5.0 kHz signal tone (4.2 and 5.946 kHz) and tones that are “distant” neighbors (+/− 1.20 octaves) to the extinguished 5.0 kHz signal tone (11.5 & 2.17 kHz). All tones are readily discriminable from the extinguished signal tone, as the threshold for discrimination is approximately 3-6% ΔF in rodents (6% ΔF range with respect to 5.0 kHZ: 4.7-5.3 kHz) (Heffner & Masterton, 1980; Talwar & Gerstein, 1998; Syka et al., 1996; Chen, Krueger-Burg, & de Hoz, 2019). Each tone frequency was presented a total 12 times. The session was divided into four continuous blocks, with 3 presentations of each tone per block in a pseudo-random order. No responses were reinforced.

Behavioral procedures for the additional subjects used in the control task are described in detail in Rotondo & Bieszczad (2020). Briefly, rats learned to press a bar for reward as described above. Subsequently, rats underwent ***single-tone sound-reward training*** in which they learned only to associate the 5.0 kHz signal tone with reward. Rats were trained to criteria (at least 70% of responses occurring to the signal tone vs. silence over for at least two consecutive days). These animals did not undergo any extinction. 48 hours following the final tone reward training session, rats received a Memory Test, as above, to determine the specificity of their behavioral responding to the signal frequency (5.0 kHz) versus four other frequencies that in this case were novel sounds (2.17 kHz, 4.2 kHz, 5.946 kHz & 11.5 kHz).

### 2.3 Pharmacological Inhibition of HDAC3 Activity

A pharmacological HDAC3 inhibitor RGFP966 was used to alter molecular mechanisms of auditory memory formation induced by learning (Bieszczad et al., 2015). Rats in this experiment were assigned to either the RGFP966 (n=7) or vehicle (n=7) condition prior to extinction training such that group performance was matched for tone-reward training. Rats received immediate post-session injections after each of the 2 extinction training sessions of RGFP966 (10 mg/kg, s.c.) or vehicle (equated for volume) (dose established [Malvaez et al., 2013], and confirmed in auditory system function [Bieszczad et al., 2015]). Post-training pharmacological treatment confines manipulation to the memory consolidation period, while avoiding potential performance effects based on perception, motivation or within-session learning.

Rats used in the control single-tone reward training task also received injections of either RGFP966 or vehicle at the dosage described above. These rats received post-session injections immediately following each of 3 single tone-reward training sessions (on days 2-4). Rats received saline injections (equated for volume) immediately following the remainder of the training sessions.

### 2.4 Auditory Brainstem Response Recording Procedure

Auditory brainstem responses (ABRs) were recorded three times in anesthetized rats (sodium pentobarbital, 50 mg/kg, i.p.) to determine learning-induced changes in subcortical sound processing: (1) 24 hours prior to tone-reward training, (2) 24 hours prior to extinction training, and (3) 24 hours following the final extinction training session. All recordings were made in a recording chamber completely separate from the training chamber and while the animal was anesthetized, which is a completely different state and context than that used in training. Stimulus presentation and neural response recordings were carried out using BioSig RZ software (TDT Inc.). Evoked potentials were recorded using a three-electrode configuration, with subdermal needle electrodes (1 kΩ) positioned at the midline along the head (recording), immediately below the left pinna (reference), and the midline on the back of the neck (ground). Sound stimuli were 60 dB SPL, 5ms pure-tones (2 ms cosine-gated rise/fall time) presented at 21 Hz to the left ear from a speaker positioned 4 cm away. Three tone frequencies (11.5, 5.946, and 5.0 kHz) were presented in a blocked format (512 stimuli per block). The averaged evoked response was used for analysis of the first positive peak (PW1) and fifth positive peak (PW5) of the waveform. Custom Matlab© scripts were used to identify peaks within the waveform and derive the trough-to-peak amplitude (μV) and peak slope (amplitude (μV)/time span (s)).

For ABR peaks PW1 and PW5, amplitude and peak slope were analyzed to determine learning-induced changes for the following groups: drug treatment groups (RGFP966-treated *vs*. vehicle-treated animals) and memory phenotype groups (frequency-specific *vs*. frequency-general memory animals). Learning induced changes in peak features were calculated as: ((Post-training amplitude – pre-training amplitude)/pre-training amplitude))*100 and ((Post-training slope – pre-training slope)/pre-training slope))*100. To determine subcortical plasticity induced by multi-tone reward training, the change in ABR peak amplitude and width were calculated between ABRs recorded 24 hours prior to the first multi-tone reward training session and ABRs recorded 24 hours following the final multi-tone reward training session. To determine subcortical plasticity induced by single-tone extinction training, the change in ABR peak amplitude and width will be determined between ABRs recorded 24 hours prior to the first extinction training session and ABRs recorded 24 hours following the final extinction training session.

ABRs recorded from rats used in the control single-tone reward training task were obtained as described above. ABRs were recorded at two time points: (1) 24 hours prior to the first single tone-reward training session and (2) 24 hours following the final tone-reward training session.

### 2.5 Auditory Cortical Recording Procedures

To determine changes in the frequency-specificity of auditory cortical bandwidth tuning, electrophysiological recordings were obtained from anesthetized subjects (sodium pentobarbital, 50mg/kg, i.p.) in an acute, terminal recording session 24-48 hours following the Extinction Memory Test. All recordings were in the same recording chamber as what was used to obtain ABRs, which was completely separate from the training chamber while the animals were in a completely different state and context than that used in training. Recordings were also obtained from a group of control rats that received multi-tone reward training, but did not undergo extinction training. All recordings were performed inside a double-walled, sound attenuated room using a linear array (1 x 4) of parylene-coated microelectrodes (1-2 MΩ, 250 μm apart) targeted to the middle cortical layers (III-IV, 400-600 μm orthogonal to the cortical surface) of the right auditory cortex. Multiple penetrations were performed across the cortical surface (M=63.55 sites/animal, SE=3.62). Acoustic stimuli were presented to the left ear from a speaker positioned ~10 cm from the ear. Sounds were 50 ms pure tones (1-9 ms cosine-gated rise/fall time) presented in a pseudorandom order (0.5-54.0 kHz in quarter-octave steps; 0-70 dB SPL in 10 dB steps; 5 repetitions) with a variable inter-stimulus interval an average of 700 +/− 100ms apart. Neural activity was amplified x1000 and digitized for subsequent off-line spike detection and analysis using custom Matlab© scripts. Recordings were bandpass filtered (0.3-3.0 kHz). Multiunit discharges were characterized using previously reported temporal and amplitude criteria (Elias et al., 2015). Acceptable spikes were designated as waveforms with peaks separated by no more than 0.6 ms and with a threshold amplitude greater than 1.5 (for the positive peak) and less than 2.0 (for the negative peak) x RMS of 500 random traces from the same recording on the same microelectrode for each site. For each recording site, tone-evoked spike rate (spikes/s) were calculated by subtracting spontaneous spiking (40 ms window prior to tone onset) from evoked-spiking within a 40 ms response-onset window (6-46 ms after each tone onset). Responses greater than +/−1.0 SEM of the spontaneous spike rate were considered true evoked responses. Tone-evoked activity was used to construct frequency-response areas (FRAs) for each recording site, which reveal the mean sound-evoked activity to each frequency/sound level combination. The borders of each FRA were determined based on a threshold firing rate value determined by its spontaneous activity. Only evoked responses greater than the mean of pre-onset spontaneous activity were considered true sound-evoked responses. The outside border of each FRA was used to determine (1) response threshold, or the lowest sound level (dB SPL) that evokes a response, (2) characteristic frequency (CF), or the frequency to which the site responds most strongly (in spikes/s) at threshold sound level, and (3) tuning bandwidth, or the breadth of frequency responsivity (in octaves) as a function of dB above response threshold. Thus, BW10, BW20, BW30, and BW40 denote bandwidth 10, 20, 30, and 40 dB SPL above threshold sound level, respectively. In order to determine tuning plasticity as a function of acoustic frequency, bandwidth data was sorted by CF to create three frequency bins: (1) sites tuned near (within +/− 1/3 octave) the extinguished 5.0 kHz signal tone frequency (4.2-6.0 kHz), (2) sites tuned near (within +/− 1/3 octave) the non-extinguished 2.17 kHz tone (1.75-2.75 kHz), and (3) sites tuned near (within +/− 1/3 octave) the non-extinguished 11.5 kHz tone (9.1-14.5 kHz).

### 2.6 Experimental Design and Statistical Analysis

Group sizes are consistent with prior reports showing brain-behavior relationships in rodent models of auditory memory (Bieszczad & Weinberger, 2010) and were sufficient to detect significant differences between treatment and untreated groups that were both frequency-specific and in the predicted direction (e.g., increased memory specificity). Furthermore, we reasoned that these data are sufficiently powered as the findings are consistent with prior findings using the HDAC3-inhibitor (RGFP966) in Bieszczad et al. (2015), Shang et al. (2019), and Bieszczad & Rotondo (2020). Measures of the plasticity of the auditory brainstem response are within-subject, providing additional power to detect significant differences. Finally, correlative data between measures of neural plasticity and learned behavior further validate the current findings. All statistical analyses were performed using SPSS software (IBM, Armonk NY).

#### 2.6.1 Behavioral Statistical Analysis

Fourteen adult male Sprague Dawley rats (RGFP966 n= 7; vehicle n = 7) were used in the extinction experiment. A two-way ANOVA (treatment group x training session *or* memory phenotype x extinction session) was used to compare group performance on the first two tone-reward training sessions and the final two tone-reward training sessions. Performance data from tone-reward training was be used to create matched groups prior to pharmacological treatment during extinction training.

There were 10 rewarded catch trials (2 per signal tone frequency in the *multiple tone detection task*) during the first day of extinction training. A mixed-model ANOVA (treatment group x frequency) was used to determine whether rats showed a flat behavioral response gradient (number of bar-presses) across acoustic frequency immediately prior to single-tone extinction training. For extinction trials during extinction training, an independent samples t-test between treatment groups *or* memory phenotype groups was used to determine differences in the number of responses to the first 15 trials during extinction session 1. In addition, an independent samples t-test between treatment groups *or* memory phenotype groups was used to test for differences in the number of responses to the first 15 trials during extinction session 2 (expressed as a proportion of responses on extinction day 1).

For the Extinction Memory Test, the distribution of bar presses among the test tone frequencies was used to determine the shape of the frequency generalization gradient. To quantify memory specificity for the extinguished signal tone, contrast measures of relative to response to the signal tone, vs. non-extinguished tones, were calculated as follows: (1) Percent of responses to extinguished signal tone – (average percent of response to non-extinguished distant tones) and (2) Percent of responses to extinguished signal tone – (average percent of responses to non-extinguished nearby tones). Negative values indicate *less* responding to the extinguished signal tone than non-extinguished tones. Single-sample t-tests were used to determine whether contrast scores were significantly different than 0.

To determine memory phenotype based on response during the Extinction Memory Test, we defined animals with frequency-specific extinction memory as those who had *negative* contrast values for both distant and nearby tones, indicating *less* bar pressing to the extinguished signal tone vs. non-extinguished signal tones (as in Rotondo & Bieszczad, 2020). A binomial test was used to determine the categorical frequency of memory phenotype by treatment condition.

#### 2.6.2 Auditory Brainstem Response Recordings Statistical Analysis

ABRs were recorded at three time points for each of the 14 RGFP966- and vehicle-treated rats in the extinction experiment (as specified in *2.4 Auditory Brainstem Response Recording Procedure*). PW5 was not able to be detected for one subject in pre-training ABRs evoked by 5.0 and 5.946 kHz. In the same subject, PW5 was not able to be detected in post-extinction ABRs evoked by 5.0 kHz. Samples sizes used all ABR analyses are specified in Tables 1–9.

**Table 1.** Changes in PW1 following multi-tone reward training. This table presents change in PW1 slope and amplitude (M ± SE) from pre- to post-multi-tone reward training. Sample sizes are in parentheses. Note that drug or vehicle treatment had not been administered at this time point, nor had general or specific extinction memory phenotype been determined.

|  |  | all | vehicle | RGFP966 | general | specific |
| --- | --- | --- | --- | --- | --- | --- |
| 5.0 kHz | amplitude | $3.78 \pm 4.01$<br>(14) | $5.13 \pm 6.34$<br>(7) | $3.24 \pm 4.46$<br>(7) | $4.46 \pm 5.18$<br>(6) | $3.98 \pm 5.56$<br>(8) |
| | slope | $2.15 \pm 3.61$<br>(14) | $2.76 \pm 5.53$<br>(7) | $2.14 \pm 4.27$<br>(7) | $3.90 \pm 6.01$<br>(6) | $1.36 \pm 4.11$<br>(8) |
| 5.946 kHz | amplitude | $1.27 \pm 4.09$<br>(14) | $5.25 \pm 5.48$<br>(7) | $-1.29 \pm 5.53$<br>(7) | $1.04 \pm 1.67$<br>(6) | $2.69 \pm 4.46$<br>(8) |
| | slope | $0.53 \pm 3.39$<br>(14) | $0.11 \pm 5.40$<br>(7) | $2.13 \pm 3.73$<br>(7) | $1.67 \pm 5.34$<br>(6) | $0.71 \pm 4.22$<br>(8) |
| 11.5 kHz | amplitude | $2.95 \pm 3.20$<br>(14) | $3.51 \pm 4.98$<br>(7) | $3.22 \pm 3.75$<br>(7) | $2.11 \pm 5.67$<br>(6) | $4.31 \pm 3.42$<br>(8) |
| | slope | $1.61 \pm 3.50$<br>(14) | $0.66 \pm 5.70$<br>(7) | $3.79 \pm 3.74$<br>(7) | $2.93 \pm 6.27$<br>(6) | $1.68 \pm 3.78$<br>(8) |

For each pair of ABRs (i.e. pre and post), data were analyzed using a mixed-model ANOVA with the within-subjects factor of *acoustic frequency* and the between-subjects factor of either *drug treatment* or *memory phenotype*. Holm-Bonferroni corrected t-tests were used to follow up significant findings where appropriate. Because this ANOVA detects differences *between* pairs of frequencies, single sample t-tests were additionally used to determine whether ABRs evoked by a single frequency showed a significant change (i.e. different from 0) over the training interval.

To limit the number of statistical comparisons, the following statistical choices were made based on *a priori* hypothesis-driven experimental goals to determine effects with respect to signal tone responses. Therefore: (1) All contrast comparisons are made with respect to 5.0 kHz, which was the extinguished signal tone. Therefore, contrasts between 5.946 and 11.5 kHz were not tested. (2) All comparisons, with the exception of the repeated-measures ANOVA, were made within (rather than between) drug-treatment groups or extinction memory phenotype groups. The goal was to determine whether a group shows signal-specific plasticity (i.e. whether there are relative differences in the representation of frequencies), rather than to compare the magnitude of plasticity across groups.

For each of the 11 rats used in the control single-tone reward training task, ABRs were recorded at two time points (as specified in *2.4 Auditory Brainstem Response Recording Procedure*). We were unable to obtain valid post-training data for 2 of 11 subjects. Sample sizes are analysis are reported in Table 5.

#### 2.6.3 Auditory Cortical Recordings Statistical Analysis

For group analysis of bandwidth, individual recording sites were treated as individual observations (No Extinction Control: n=6 subjects/37 recordings sites near 5.0 kHz/30 recording sites near 2.17 kHz kHz/57 sites near 11.5 kHz; vehicle: n=7 subjects/27recordings sites near 5.0 kHz/22 recording sites near 2.17 kHz kHz/37 sites near 11.5 kHz; RGFP966: n=7 subjects/23 recordings sites near 5.0 kHz/27 recording sites near 2.17 kHz kHz/34 sites near 11.5 kHz). In 3 subjects (all drug-treated individuals with specific-memory phenotype), no cortical sites tuned near 5.0 kHz were detected and therefore these subjects are excluded from those analyses. Sample sizes for bandwidth analysis are reported in Tables 10 and 11. Differences in tuning bandwidth within a frequency bin was compared among conditions (vehicle, RGFP966, and No Extinction Control) using one-way ANOVA. Pairwise comparisons were made with Holm-Bonferroni corrected two-tailed t-tests. Corrected p-values are reported.

#### 2.6.4 Correlative Data

To limit the number of statistical comparisons, we set the *a priori* criterion that brain-behavior Pearson correlations would be pursued only for brain measures that showed learning-induced plasticity. The number of subjects available for each correlation (determined by subject attrition in neural measures, as described in sections *2.6.2 Auditory Brainstem Response Recordings Statistical Analysis* and *2.6.3 Auditory Cortical Recordings Statistical Analysis*) is detailed in Table 12.

Measures used for Pearson correlations are as follows. For behavioral measures, contrast measures were derived from relative number of responses to the extinguished 5.0 kHz tone *vs*. non-extinguished tones, including (1) the difference in number of bar-presses to 5.0 kHz vs. all non-extinguished tones, (2) the difference in number of bar-presses to 5.0 kHz vs. 5.946 kHz, and (3) the difference in number of bar-presses to 5.0 kHz vs. 11.5 kHz. For ABR data, contrast measures for the changes in ABRs evoked by pairs of frequencies were calculated for each subject, including (1) the difference in amplitude or slope change between ABRs evoked by 5.0 vs. 5.946 kHz, (2) the difference in amplitude or slope change between ABRs evoked by 5.0 vs. 5.946 kHz, and (3) the average difference in amplitude or slope between ABRs evoked by 5.0 vs 5.946 and 5.0 vs. 11.5 kHz. For auditory cortical tuning bandwidth, an average bandwidth score for BW20 was computed for each individual.

## 3 Results

### 3.1 HDAC3 Inhibition Promotes Frequency-Specific Extinction Memory

Multi-tone reward training catch trials were used prior to extinction training to determine whether there was a flat response gradient for each subject across the frequency dimension. All animals pressed equivalently to each of the five training frequencies during the rewarded catch trials in the initial portion of the first extinction session. A mixed-model ANOVA revealed no main effect for frequency (F(4,48) = 1.531, p = 0.208, no main effect for drug treatment group (F(1,12) = 0.001, p = 0.976, and no *frequency x drug treatment group* interaction (F (4,48) = 2.053, p = 0.104) (**Vehicle:** 2.17 kHz-M = 4.14, SE = 0.40; 4.0 kHz-M = 4.00, SE = 0.21; 5.0 kHz-M = 3.85, SE = 0.26; 5.946 kHz-M = 4.00, SE = 0.21; 11.5 kHz-M = 3.85, SE = 0.33; **RGFP966:** M = 3.57, SE = 0.81; 4.0 kHz-M = 3.85, SE = 0.81; 5.0 kHz-M = 4.28, SE = 1.10; 5.946 kHz-M = 4.42, SE = 1.10; 11.5 kHz-M = 3.57, SE = 0.80)

Following multi-tone reward training and prior to extinction training, rats were split into two performance-matched groups (Fig. 1a) (*number of training days:* RGFP966: M= 8.142, SE = 0.961, VEH: M = 8.285, SE = 0.644, independent samples t-test: t(12) = 0.123, p = 0.903; *first and final performance:* RGFP966: first two training sessions-M=0.096, SE=0.027, last two training sessions: M=0.796, SE=0.0233; VEH: first two training sessions-M=0.111, SE=0.023, last two training sessions: M=0.771, SE=0.035; Two-way ANOVA: treatment group-F(1,24)= 0.34, p = 0.855; treatment group *x* session-F(1,24)=0.494, p=0.489).

**Figure 1.**
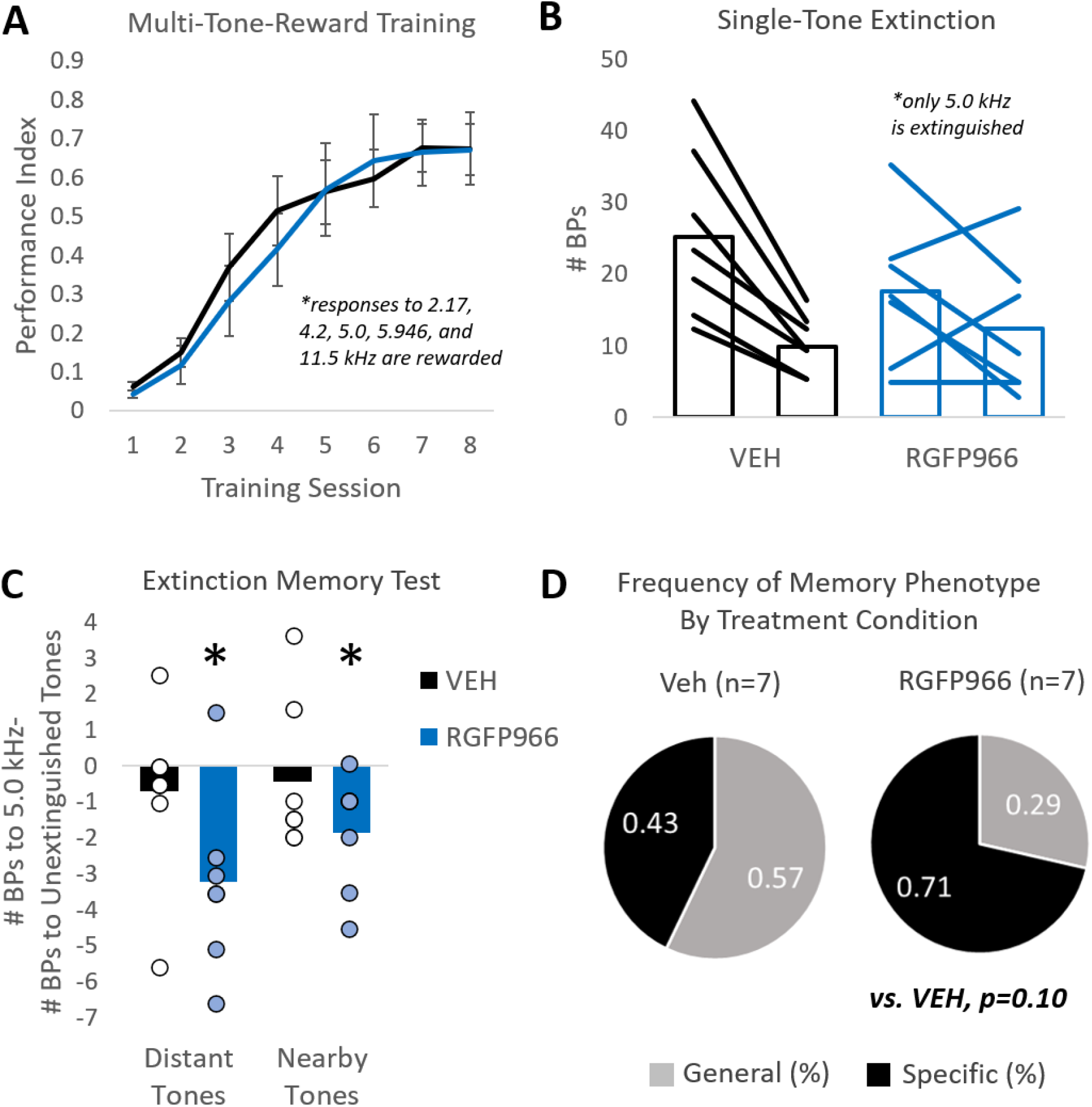
Post-training treatment with HDAC3 inhibitor RGF966 promotes frequency-specific extinction memory. **(A)** RGFP966-treated rats (n=7) learned to perform the multi-tone reward training task to an equal level of final performance as vehicle-treated rats (n=7). Five different tone frequencies are presented; responses to all frequencies are rewarded. Bar-presses to tones were rewarded while bar-presses during silent inter-tone/trial-intervals were unrewarded and signaled as errors with a time-out and flashing house-light. **(B)** During single-tone extinction training, only 5.0 kHz is presented. No responses are reinforced. Bars represent that average number of bar presses made by vehicle and RGFP966-treated animals on extinction day 1 and 2. Lines represent behavioral trajectory of individual subjects. **(C)** Quantifying the shape of the response distribution using relative measures of responding to the signal tone vs. other test tone frequencies reveals that RGFP966-treated animals behaviorally discriminate. They respond to the signal tone more than both distant (2.17 and 11.5 kHz) tones (*left*) and nearby (4.2 and 5.946 kHz) tones (*right*). Vehicle-treated rats do not discriminate, responding equally to signal tone vs. nearby or distant tones (n=7). **(D)** RGFP966 treatment increases the proportion of individuals with frequency-specific extinction memory type, compared with vehicle-treated individuals. *p<0.05

Following each of two single-tone extinction training sessions, rats received injections of the HDAC3 inhibitor, RGFP966 (Abcam Inc., 10 mg/kg, n=7) or vehicle (equated for volume; n=7). During the first session of single-tone extinction training (Fig. 1b), prior to pharmacological treatment, there were no group differences in the number of bar presses made to the 5.0 kHz extinction signal during the 1^st^ 15 extinction trials (RGFP966: M = 17.57, SE = 3.62; VEH: M = 25.28, SE = 4.48; independent samples t-test: t(12) = 1.312, p=0.214). During the second session of single-tone extinction training, there were no group differences in the number of bar presses made to 5.0 kHz extinction signal, expressed as a proportion of bar presses made during extinction session 1 (RGFP966: M = 0.88, SE = 0.29; VEH: M = 0.40, SE = 0.02; independent samples t-test: t(12) = −1.621, p = 0.1308), though RGFP966-treated animals exhibited a less consistent response trajectory from one session to the second. Nevertheless, HDAC3 inhibition does not significantly alter the rate of extinction learning over days.

However, during the Extinction Memory Test (Fig. 1c), RGFP966-treated rats exhibited a frequency-specific behavioral response distribution, which characterizes a “specific” extinction memory phenotype. RGFP966-treated animals made significantly more bar presses to the non-extinguished distant tones (M= −3.21, SE = 0.93; single sample t-test: t(6) = −3.43, p = 0.013) and nearby tones (M= −1.85, SE = 0.60; single sample t-test: t(6) = −3.07, p = 0.021) than the extinguished 5.0 kHz signal tone. In contrast, vehicle-treated animals did not behaviorally discriminate the extinguished signal tone from either nearby tones (M = −0.42, SE = 0.79; single sample t-test: t(6) = −0.537, p = 0.610) or distant tones (M = −0.71, SE = 0.90; single sample t-test: t(6) = −0.788, p = 0.460), which characterizes a “general” extinction memory phenotype. There were no group differences in the number of bar presses to tones during the Extinction Memory Test (RGFP966: M = 12.28, SE = 1.89; VEH: M = 8.71, SE = 2.83; independent samples t-test: t(12) = −1.046, p = 0.316), suggesting that the primary effect of RGFP966 treatment on extinction memory was to shift within-group variability to frequency specific memory. Indeed, compared to vehicle treatment, there was a trend for RGFP955 treatment to increase the proportion of individuals the group with “specific” memory (Fig 1d; Vehicle: n = 3/7; RGFP966: n = 5/7; binomial test: p = 0.100). These findings are consistent with previous work demonstrating that HDAC3 is a mechanism that drives memory towards specificity (Bieszczad et al., 2015; Shang et al., 2019; Rotondo & Bieszczad, 2020). These findings for the first time extend this role to *extinction* memory.

To later determine whether the neural correlates of frequency-specific learning depended on drug (vs. vehicle) treatment, the data were also analyzed re-grouped by extinction memory phenotype (frequency-specific or frequency-general), rather than treatment group. There were no group differences in performance during multi-tone reward training (*number of training days:* General: M= 9.333, SE = 0.988, Specific: M = 8.250, SE = 0.674, independent samples t-test: t(12) = 0.938, p = 0.366; *first and final performance:* General: First two training sessions-M=0.092, SE=0.031, Last two training session: 0.797, SE=0.029; Specific: First two training sessions: M=0.112, SE=0.020, Last two training sessions: M=0.773, SE=0.029; Two-way ANOVA: treatment group-F(1,24)= 0.004, p = 0.953; treatment group *x* session-F(1,24)=0.642, p=0.431). During the first session of single-tone extinction training, there were no group differences in the number of bar presses made to the 5.0 kHz extinction signal during the 1^st^ 15 extinction trials (Specific: M=22.75, SE=4.109; General: M=19.667, SE=4.765; independent samples t-test: t(12) = − 0.490, p=0.632). During the second session of single-tone extinction training, there were no group differences in the number of bar presses made to 5.0 kHz extinction signal, expressed as a proportion of bar presses made during extinction session 1 (Specific: M=0.638, SE=0.258; General: M=0.716, SE=0.164; independent samples t-test: t(12) = 0.236, p=0.817. During the Extinction Memory Test, rats with frequency-specific extinction memory made significantly more bar presses to the non-extinguished distant tones (M= −2.937, SE = 0.770; single sample t-test: t(7) = −3.814, p = 0.006) and nearby tones (M= −1.625, SE = 0.309; single sample t-test: t(7) = −5.245, p = 0.001) than the extinguished 5.0 kHz signal tone. In contrast, rats with frequency-general memories did not behaviorally discriminate the extinguished signal tone from either nearby tones (M = −0.500, SE = 1.157; single sample t-test: t(5) = −0.435, p = 0.599) or distant tones (M = −0.667, SE = 1.187; single sample t-test: t(5) = −0.435, p = 0.681). There were no group differences in the number of bar presses to tones during the Extinction Memory Test (Specific: M = 9.75, SE = 2.00; General: M = 11.5, SE = 3.149; independent samples t-test: t(12) = 0.490, p = 0.632), suggesting that the primary difference between these groups was the frequency-specificity of extinction memory.

### 3.2 Multi-Tone Reward Training Results In Subcortical Plasticity In The Processing Of Frequencies Associated With Reward

Previous work has used auditory brainstem responses (ABRs) to demonstrate learning-induced subcortical plasticity in the representation of behaviorally relevant sounds after sound-reward learning (Bieszczad & Rotondo, 2020). The present study also investigated changes in auditory brainstem responses evoked by a pure tones during the initial phase of training. To do so, ABRs were compared pre- and post-multi-tone reward training (i.e., prior to any extinction training). Specifically, the first positive peak (PW1) and the fifth positive peak (PW5) were analyzed for changes in amplitude and slope. These peaks were chosen because they represent distinct levels of subcortical auditory processing. PW1 represents potentials generated at approximately the level of the auditory nerve (Starr, 1976; Møller & Jannetta, 1985; Chen & Chen, 1991; Hall, 2007), and PW5 likely represents potentials generated by multiple nuclei within the auditory midbrain, including the nuclei of the lateral lemniscus and the inferior colliculus (Starr, 1976; Møller & Jannetta, 1985; Chen & Chen, 1991; Hall, 2007).

Here, multi-tone reward training was used to generate a flat behavioral gradient with respect to sound frequency to set the stage for the ability to detect frequency-specific effects after extinction. Because all frequencies had identical behavioral relevance at this stage, one of two possible outcomes were predicted for the neural correlates of multi-tone reward training. One prediction was that subcortical processing of frequency would not change from pre- to post-training if subjects are not required, or do not learn to use acoustic frequency as the relevant dimension to cue behavior, which has been shown in prior work (Bieszczad & Rotondo, 2020). An alternate prediction is that the subcortical processing of frequency *would* change from pre- to post- training, but the form and magnitude of subcortical changes would not differ among different frequencies within the same individual; the changes might be a general change in sound-processing.

In support of the second prediction, an increase in PW5 amplitude and slope was significant or trending toward significance in ABRs evoked by all frequencies presented (i.e., the same frequencies that had been predicted reward during training) (Fig. 2b; Table 3). Importantly, a mixed-model ANOVA revealed no main effect for acoustic frequency for peak amplitude (F(2,22) = 0.081, p = 0.923) or peak slope (F(2,22) = 0.107, p = 0.899), indicating that the magnitude of change across frequencies was not significantly different (Fig. 3b; Table 4). This indicates the change was not frequency-specific, but a *general* learning-induced change in the subcortical processing of sound. To determine whether PW5 could be at all sensitive to frequency-specific effects, we analyzed ABRs in the same way from animals trained in the control single-tone sound-reward training task. Similarly, this task does not require that animals learn about acoustic frequency *per se*. In this case, animals that did develop a frequency-specific memory phenotype, including the RGFP966-treated animals, exhibited a signal-specific increase in PW5 amplitude and slope in ABRs evoked by 5.0 kHz (Table 5). In contrast, animals that formed frequency-general memory, including vehicle-treated animals, did not show any significant changes in PW5 amplitude or slope in ABRs evoked by any frequency. Altogether, these data suggest that learning-induced changes to sound-evoked PW5 amplitude and slope are sensitive to the strategy animals use to learn the behavioral significance of sounds.

**Figure 2.**
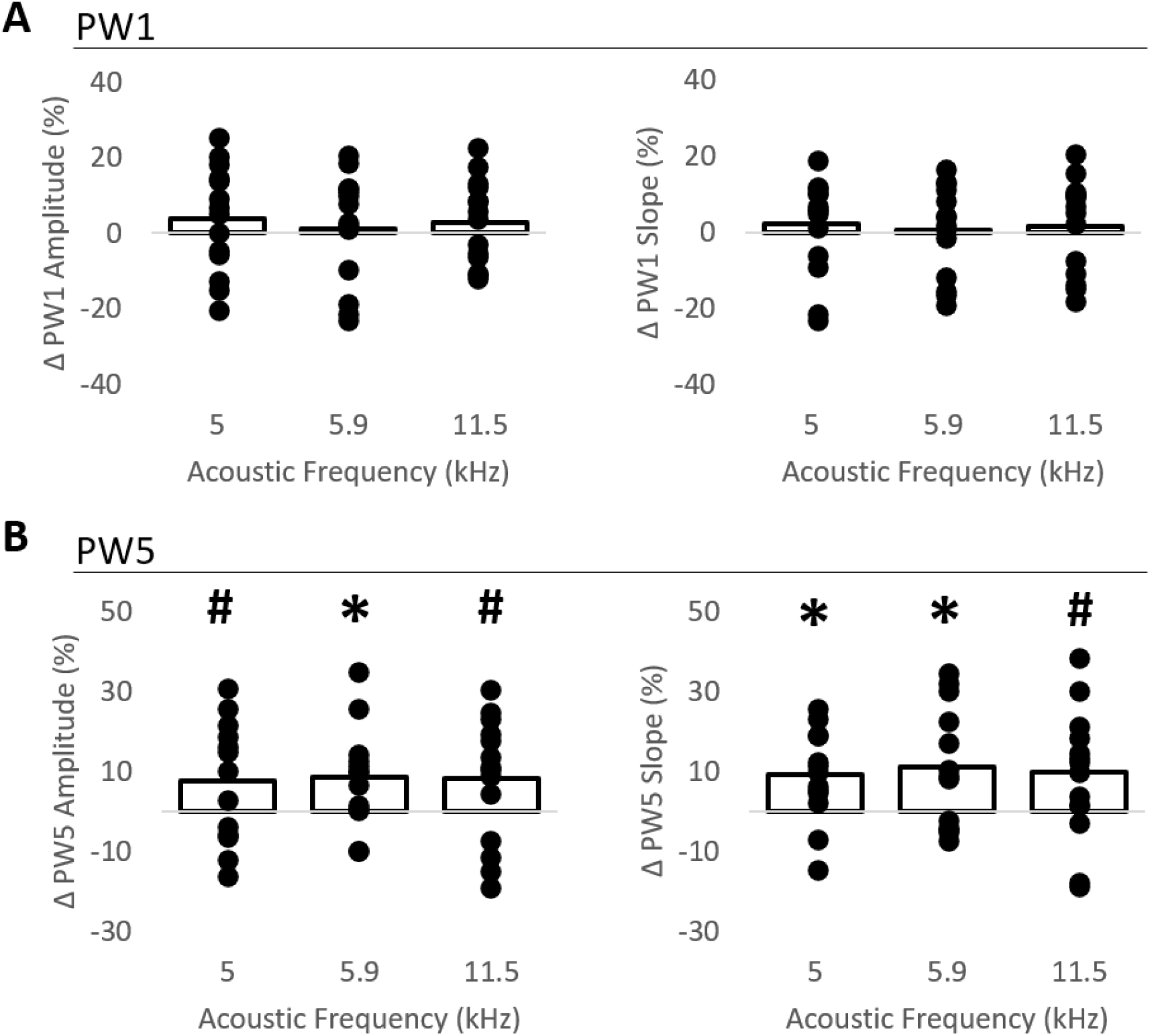
Multi-tone reward training is associated with increased amplitude and slope in PW5 of the ABR. **(A)** There was no change from pre- to post- multi-tone reward training in the amplitude (left) or slope (right) of PW1 in ABRs evoked by a subset of the tones that had been associated with reward (5.0, 5.946, or 11.5 kHz). **(B)** There was a significant increase or a trend toward a significant increase from pre- post- multi- tone reward training in the amplitude (left) and slope (right) of PW5 in ABRs evoked the 5.0, 5.946, and 11.5 kHz tones. Bars indicate average change. Dots indicate individual data points. *p<0.05, ^#^p<0.10

**Figure 3.**
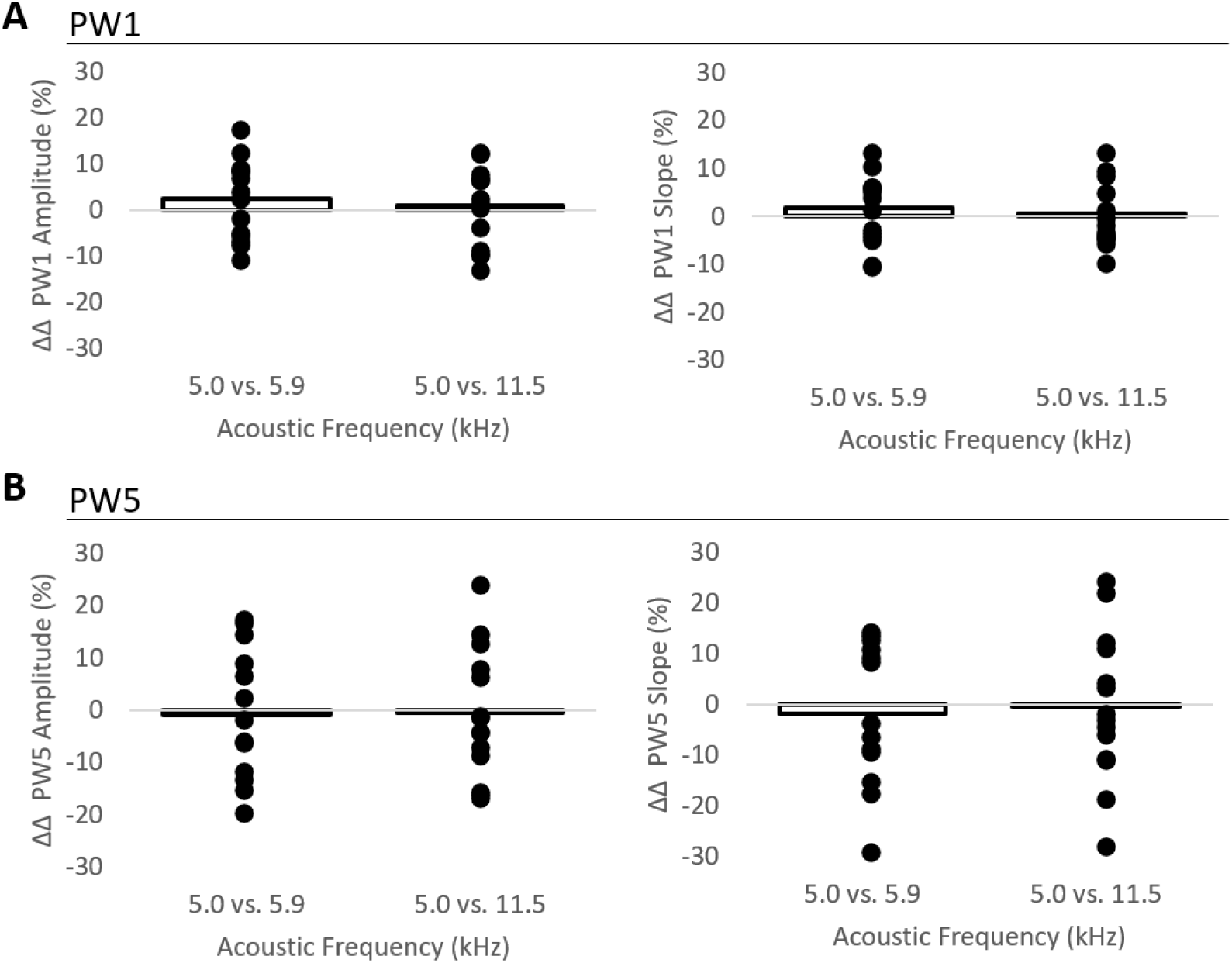
Multi-tone reward training does not produce different changes in the amplitude or slope of ABRs evoked by different frequencies. The relative change in the amplitude (left) or slope (right) of **(A)** PW1 or **(B)** PW5 was not different among ABRs evoked by different frequencies. Bars indicate average change. Dots indicate individual data points.

Interestingly, no significant change was observed for PW1 amplitude or slope for any of the frequencies presented (Fig. 2a; Table 1). A mixed**-** model ANOVA further revealed no main effect for acoustic frequency for peak amplitude (F(2,24) = 0.460, p = 0.637) or peak slope (F(2,24) = 0.312, p = 0.735) (Fig. 3a; Table 2). Therefore, a dissociation is observed in learning-induced plasticity between more peripheral (i.e., PW1) versus midbrain-generated (i.e., PW5) auditory responses in response to learning about multiple frequencies. Of note, prior findings have shown that PW1 is subject to learning-induced plasticity as it can change in animals that learn with a frequency-specific strategy (Rotondo & Bieszczad 2020). However, unlike PW5, the learning-induced changes in PW1 do not appear to be sensitive to learning that generalizes across frequency, which was the case in this multi-tone task.

**Table 2.**
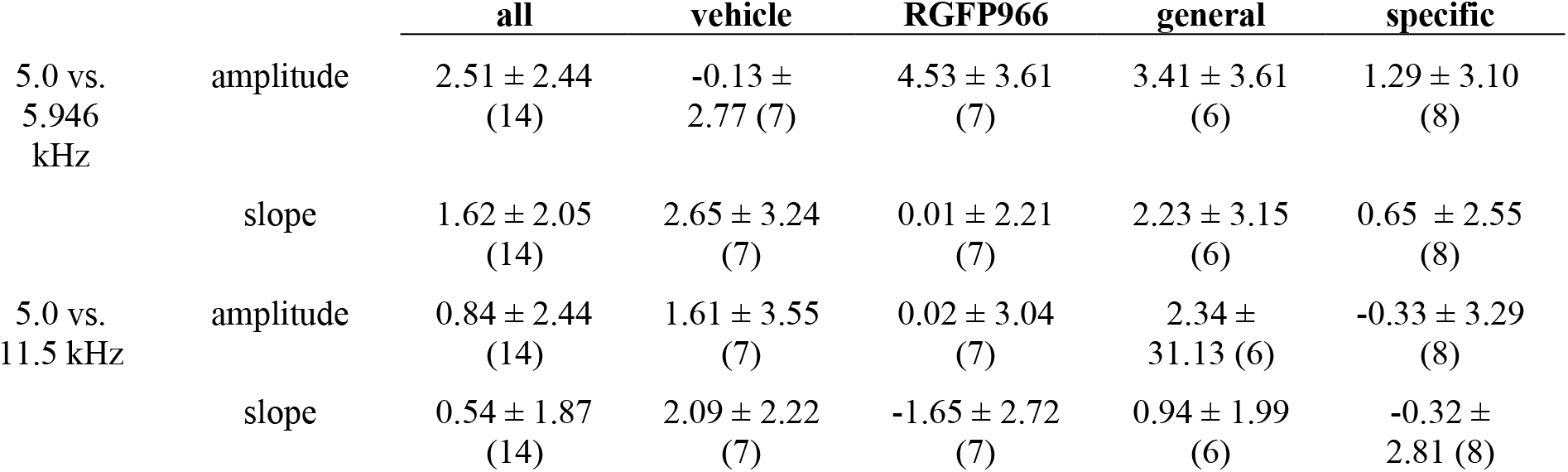
Relative changes in PW1 between pairs of frequencies following multi-tone reward training. This table presents the relative difference in change in PW1 slope and amplitude (M ± SE) between pairs of frequencies from pre- to post-multi-tone reward training. Sample sizes are in parentheses. Note that drug or vehicle treatment had not been administered at this time point, nor had general or specific extinction memory phenotype been determined.

**Table 3.** Changes in PW5 following multi-tone reward training. This table presents change in PW5 slope and amplitude (M ± SE) from pre- to post-multi-tone reward training. Sample sizes are in parentheses. Note that drug or vehicle treatment had not been administered at this time point, nor had general or specific extinction memory phenotype been determined. *p<0.05, **p<0.01, ^#^p<0.10

|  |  | all | vehicle | RGFP966 | general | specific |
| --- | --- | --- | --- | --- | --- | --- |
| 5.0 kHz | amplitude | 7.51 ± 4.20<sup>#1</sup><br>(13) | 9.77 ± 7.08<br>(7) | 5.01 ± 5.53<br>(6) | 6.79 ± 5.84<br>(6) | 8.25 ± 6.42<br>± (7) |
| | slope | 9.26 ±<br>3.19*<sup>2</sup> (13) | 12.12 ±<br>3.02**<sup>3</sup> (7) | 5.99 ± 5.07<br>(6) | 8.97 ±<br>4.31<sup>#4</sup> (6) | 9.52 ± 4.94<br>(7) |
| 5.946 kHz | amplitude | 8.51 ±<br>3.47*<sup>5</sup> (13) | 5.10 ± 3.69<br>± (7) | 12.48 ± 6.58<br>(6) | 6.23 ± 4.86<br>(6) | 10.46 ±<br>5.16<sup>#6</sup> (7) |
| | slope | 11.06 ±<br>4.20*<sup>7</sup> (13) | 10.06 ±<br>6.51 (7) | 12.22 ± 6.54<br>(6) | 11.66 ±<br>6.04 (6) | 10.54 ±<br>6.29 (7) |
| 11.5 kHz | amplitude | 8.07 ± 4.47<sup>#8</sup><br>(14) | 8.06 ± 7.63<br>(7) | 8.21 ± 6.75<br>(7) | 10.63 ±<br>5.65 (6) | 6.26 ± 6.12<br>(8) |
| | slope | 9.80 ± 4.59<sup>#9</sup><br>(14) | 15.56 ±<br>6.00 (7) | 2.91 ± 7.19<br>(7) | 11.27 ±<br>7.46 (6) | 7.71 ±<br>5.39*<sup>10</sup> (8) |

<sup>1</sup> One sample t-test: t(12) = 1.801, p = 0.096<sup>#</sup>
<sup>2</sup> One sample t-test: t(12) = 2.905, p = 0.013<sup>#</sup>
<sup>3</sup> One sample t-test: t(6) = 4.004, p = 0.007\*\*
<sup>4</sup> One sample t-test: t(5) = 2.078, p = 0.092<sup>#</sup>
<sup>5</sup> One sample t-test: t(12) = 2.447, p = 0.030\*
<sup>6</sup> One sample t-test: t(6) = 2.028, p = 0.088<sup>#</sup>
<sup>7</sup> One sample t-test: t(12) = 2.629, p = 0.022\*
<sup>8</sup> One sample t-test: t(13) = 1.805, p = 0.094<sup>#</sup>
<sup>9</sup> One sample t-test: t(13) = 2.134, p = 0.052<sup>#</sup>
<sup>10</sup> One sample t-test: t(5) = 2.592, p = 0.048\*

**Table 4.** Relative changes in PW5 between pairs of frequencies following multi-tone reward training. This table presents the relative difference in change in PW1 slope and amplitude (M ± SE) between pairs of frequencies from pre- to post-multi-tone reward training. Sample sizes are in parentheses. Note that drug or vehicle treatment had not been administered at this time point, nor had general or specific extinction memory phenotype been determined.

|  |  | all | vehicle | RGFP966 | general | specific |
| --- | --- | --- | --- | --- | --- | --- |
| 5.0 vs.<br>5.946 kHz | amplitude | 0.93 ± 3.50<br>(13) | 4.67 ± 4.44<br>(7) | -7.47 ± 4.50<br>(6) | 0.55 ± 5.42<br>(6) | -2.21 ±<br>4.90 (7) |
| | slope | -1.79 ± 3.77<br>(13) | 2.05 ± 6.55<br>(7) | -6.28 ± 3.61<br>(6) | -2.96 ±<br>7.26 (6) | -1.02 ±<br>3.86 (7) |
| 5.0 vs. 11.5<br>kHz | amplitude | -0.49 ± 3.52<br>(14) | 1.71 ± 5.63<br>(7) | -3.92 ± 3.63<br>(7) | -3.85 ±<br>5.11 (6) | 0.96 ± 4.50<br>(7) |
| | slope | -0.53 ± 4.16<br>(14) | -3.45 ±<br>6.21 (7) | 2.17 ± 5.11<br>(7) | -2.30 ±<br>7.48 (6) | 0.61 ± 4.21<br>(7) |

**Table 5.** Changes in PW5 following single-tone reward training. This table presents change in PW5 slope and amplitude (M ± SE) from pre- to post-single-tone reward training. Sample sizes are in parentheses. *p<0.05, **p<0.01, ^#^p<0.10

|  |  | vehicle | RGFP966 | general | specific |
| --- | --- | --- | --- | --- | --- |
| 5.0 kHz | amplitude | 7.14 ± 4.15<br>(7) | 43.66 ± 7.24<br>(4)**<sup>1</sup> | 2.21 ± 4.32<br>(6) | 33.00 ±<br>9.47 (5)*<sup>2</sup> |
| | slope | 4.27 ± 7.51<br>(7) | 20.35 ± 4.82<br>(4)*<sup>3</sup> | -2.77 ±<br>5.52 (6) | 25.60 ±<br>4.95 (5)**<sup>4</sup> |
| 5.946 kHz | amplitude | 7.66 ± 4.92<br>(7) | 5.30 ± 4.88<br>(4) | 4.61 ± 6.23<br>(6) | 9.42 ± 4.87<br>(5) |
| | slope | -0.70 ±<br>4.23 (7) | 0.24 ± 3.32<br>(4) | -1.21 ±<br>4.11 (6) | 2.06 ± 4.12<br>(5) |
| 11.5 kHz | amplitude | 0.56 ± 4.47<br>(7) | 1.46 ± 5.07<br>(4) | 2.40 ± 5.74<br>(6) | -0.55 ±<br>4.62 (5) |
| | slope | 1.19 ± 2.94<br>(7) | 2.97 ± 5.70<br>(4) | -0.22 ±<br>3.08 (6) | 4.03 ± 4.40<br>(5) |

<sup>1</sup> One sample t-test: t(3) = 6.02, p = 0.009\*\*
<sup>2</sup> One sample t-test: t(4) = 3.485, p = 0.025\*
<sup>3</sup> One-sample t-test: t(3) = 4.22, p = 0.0241\*
<sup>4</sup> One sample t-test: t(4) = 5.170, p = 0.006\*\*

Importantly, for both PW1 and PW5, mixed-model ANOVAs indicated no significant effects for subsequent drug treatment group for amplitude (PW1: F(1,12) = 0.184, p = 0.676; PW5: F(1,11) = 0.014, p = 0.907) or slope (PW1: F(1,12) = 0.055, p = 0.819; PW5: F(1,11) = 0.919, p = 0.358). Further, there were no significant *subsequent drug treatment group x frequency* interactions for amplitude (PW1: F(2,24) = 0.946, p = 0.402; PW5: F(2,22) = 1.545, p = 0.236) or slope (PW1: F(2,24) = 0.529, p = 0.596; PW5: F(2,22) = 0.351, p = 0.351). This indicates that, at a time point *prior to* administration of drug (vs. vehicle) treatment during the next extinction training stage, learning-induced neural plasticity was equivalent between groups.

Similarly, for both PW1 and PW5, mixed-model ANOVAs indicated no significant effects for subsequent extinction memory phenotype for amplitude (PW1: F(1,12) = 0.027, p = 0.873; PW5: F(1,11) = 0.002, p = 0.967) or slope (PW1: F(1,12) = 0.060, p = 0.811; PW5: F(1,11) = 0.033, p = 0.859). Further, there were no significant *subsequent extinction memory phenotype x frequency* interactions for amplitude (PW1: F(2,24) = 0.164, p = 0.850; PW5: F(2,22) = 0.891, p = 0.366) or for slope (PW1: F(2,24) = 0.094, p = 0.910; PW5: F(2,22) = 0.052, p = 0.890) (see table 2, 3). This suggests that prior to extinction training, subcortical plasticity due to multi-tone reward training did not have any predictive value for whether subsequent extinction memory would be a frequency-specific or -general phenotype at the group level.

### 3.3 HDAC3 Inhibition During Single-Tone Extinction Training Is Associated With Signal-Specific Subcortical Plasticity

#### 3.3.1 Changes to PW5 amplitude reverse with frequency-specificity after extinction with HDAC3-inhbition

To investigate extinction-related subcortical plasticity, ABRs were compared between the pre- and post-single-tone extinction training time points. Given that HDAC3 inhibition during single-tone extinction training promotes the development of signal-specific extinction memory revealed behaviorally, it was hypothesized that the group treated with the HDAC3 inhibitor would show matching extinction-induced subcortical plasticity that was specific to the 5.0 kHz extinction tone frequency.

Among RGFP966-treated animals, one sample t-tests revealed a significant decrease in PW5 amplitude over the course of extinction training for the 5.0 kHz extinction signal tone. This effect was specific to about one-quarter octave of the signal tone as the 5.946 kHz non-extinguished tone also showed a significant PW5 amplitude decrease, though the magnitude of the change was significantly smaller (Table 9). There were no significant changes in the 11.5 kHz non-extinguished tone (Fig. 6b, Table 8). Among vehicle-treated animals, one sample t-tests revealed a significant decrease in PW5 amplitude for the 5.0 kHz extinction signal tone and the 11.5 kHz non-extinguished tone, and a trend toward a significant amplitude decrease for the 5.946 kHz non-extinguished tone (Fig. 6a, Table 8). Analysis of the percent change of PW5 amplitude (pre- to post- extinction) between pairs of frequencies using mixed-model ANOVA revealed a significant main effect for frequency (F(2,24) = 5.457, p = 0.011), no significant effect for treatment group (F(1,12) = 1.034, p = 0.329) but—importantly—a significant *treatment group x frequency* interaction (F(2, 24) = 5.605, p = 0.010) to indicate frequency-specific changes depended on treatment (or not) with the HDAC3 inhibitor. Follow up tests revealed that among RGFP966-treated animals, there was a significant difference in the *magnitude* of PW5 amplitude change: there was a greater amplitude decrease in ABRs evoked by the 5.0 kHz extinction signal tone compared to the 5.946 kHz non-extinguished signal tone and the 11.5 kHz non-extinguished signal tone, which is interpreted as a frequency-specific effect to the 5.0 kHz signal. (Fig. 6b, Table 9). In contrast, among vehicle-treated animals, there were no differences in the magnitude of PW5 amplitude change across pairs of frequencies, which is interpreted as a generalized effect across frequency (Fig. 6a, Table 9).

Grouping the data by extinction memory phenotype revealed the same pattern of effects on the PW5. Among animals with frequency-specific extinction memory, one sample t-tests revealed a significant decrease in PW5 amplitude over the course of extinction training for the 5.0 kHz extinction signal tone and a trend toward a significant decrease for the 5.946 kHz and 11.5 kHz non-extinguished tones (Table 8). Among animals with frequency-general extinction memory, one sample t-tests revealed a significant decrease in PW5 amplitude over the course of extinction training for the 5.0 kHz extinction signal tone and for the 5.946 kHz and 11.5 kHz non-extinguished tones (Table 8). Analysis of the percent change of PW5 amplitude (pre- to post- extinction) between pairs of frequencies using mixed-model ANOVA revealed a significant main effect for frequency (F(2,24) = 5.457, p = 0.011), though no significant main effect for memory phenotype (F(1,12) = 0.086, p = 0.775). As expected, there was a significant *memory phenotype x frequency* interaction (F(2, 24) = 3.896, p = 0.034), which indicated frequency-specific neural effects that depended on whether memory was frequency specific (or not). Likewise, follow-up tests revealed that among animals with frequency-specific extinction memory, there was a significant difference in the *magnitude* of PW5 amplitude change: there was a greater amplitude decrease in ABRs evoked by the 5.0 kHz extinction signal tone compared to the 5.946 kHz non-extinguished signal tone and the 11.5 kHz non-extinguished signal tone (Table 9). In contrast, among animals with frequency-general extinction memory, there were no differences in the magnitude of PW5 amplitude change across pairs of frequencies, which indicated a generalized effect across frequency (Table 9).

Overall, there were significant changes in PW5 amplitude over the course of extinction training that occurred in the opposite direction as those due to initial sound-reward training. While changes in PW5 amplitude over the course of extinction training were *not* limited to the 5.0 kHz extinction tone frequency in any group, the magnitude of decrease was larger for the 5.0 kHz non-extinguished frequency than the non-extinguished frequencies in groups that developed frequency-specific extinction memory.

#### 3.3.2 Changes to PW5 slope maintain after extinction with HDAC3-inhbition

In contrast to PW5 amplitude, one sample t-tests revealed that the slope of PW5 did not change significantly over the course of extinction for any frequency in any drug treatment group or memory phenotype group (Fig. 7, Tables 8, 9). Among drug treatment groups, analysis of the percent change of PW5 slope between pairs of frequencies revealed no main effect for frequency (F(2,24) = 0.133, p=0.877), no main effect for drug treatment group (F(1,12) = 0.528, p = 0.568), and no *drug treatment group x frequency* interaction (F(2, 24) = 0.083, p = 0.778). Similarly, among memory phenotype groups, there was no main effect for frequency (F(2,24) = 0.133, p=0.877), no main effect for memory phenotype (F(1,12)= 0.001, p = 0.982), and no *memory phenotype x frequency* interaction (F(2,24) = 0.282, p= 0.742). Thus, in contrast to the reversal of multi-tone reward training-induced changes observed with PW5 amplitude, the slope of PW5 exhibited a maintenance of earlier learning-induced plasticity throughout extinction training.

#### 3.3.3 PW1 is not sensitive to learning-induced extinction after multi-tone sound-reward training

Among drug treatment groups, there were no changes observed for PW1 amplitude or slope for any of the frequencies presented (Figs. 4, 5; Tables 6, 7). A mixed**-** model ANOVA revealed no main effect for acoustic frequency for peak amplitude (F(2,24) = 0.417, p = 0.664) or peak slope (F(2,24) = 0.722, p = 0.496). There was no main effect for treatment group for peak amplitude (F(1,12) = 0.006, p = 0.938) or peak slope (F(1,12) = 0.063, p = 0.807) and no *drug treatment group x frequency* interactions for amplitude (F(2,24) = 0.249, p = 0.782) or slope (F(2,24) = 0.455, p = 0.640) (Figs. 2.8, 2.9; Tables 2.5, 2.6). Further, a separate mixed-model ANOVA revealed no main effect for extinction memory phenotype for peak amplitude (F(1,12) = 0.341, p = 0.570) or slope (F(1,12) = 0.362, p = 0.559) and no *extinction memory phenotype x frequency* interaction for peak amplitude (F(2,24) = 0.180, p = 0.836) or slope (F(2,24) = 0.457, p = 0.638). Therefore, neither multi-tone reward training nor single-tone extinction appeared to have consistent effects on auditory processing at this level.

**Figure 4.**
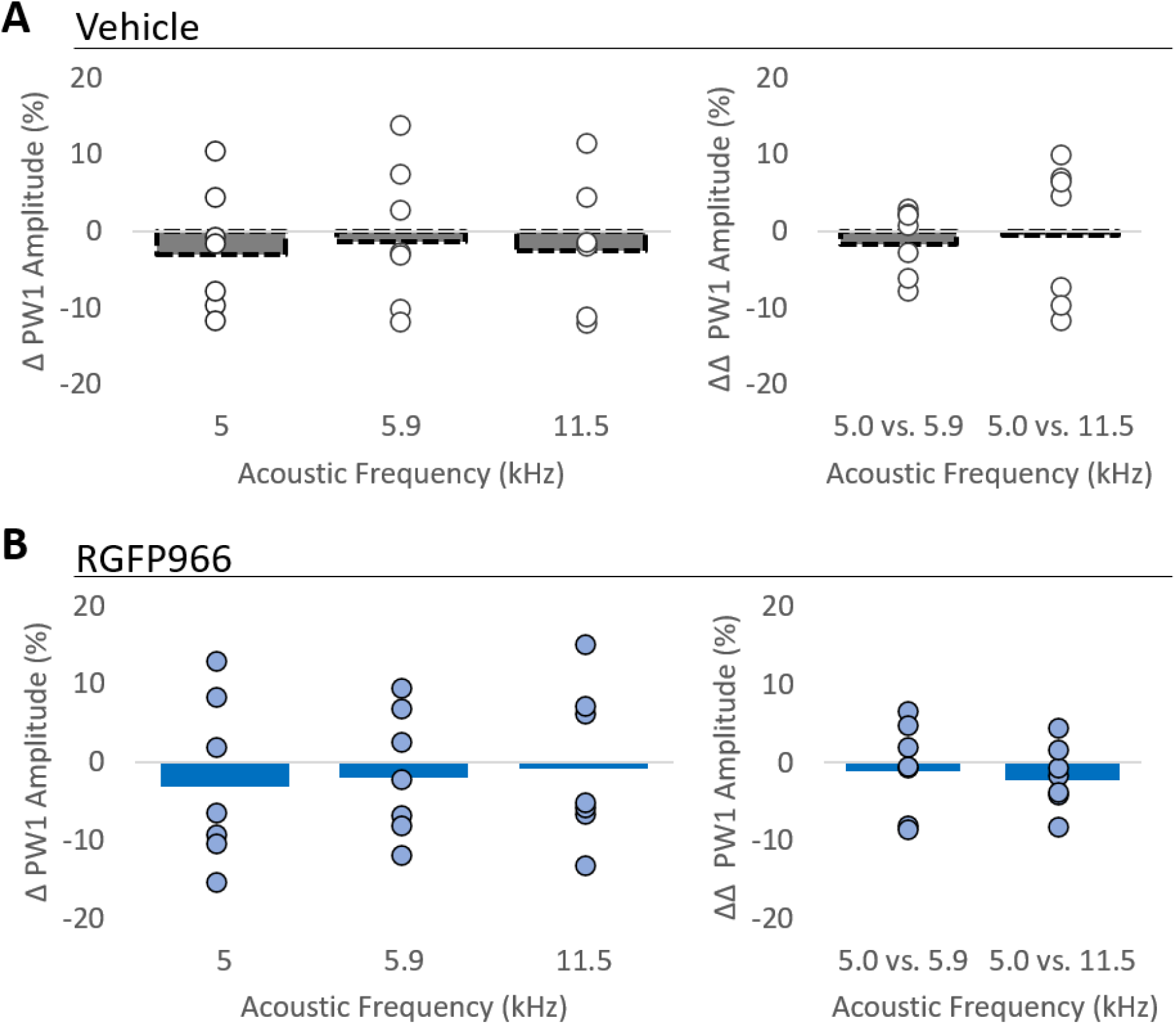
Single-tone extinction training does not result in PW1 amplitude changes in drug treatment groups. Among both (**A**) vehicle- and **(B)** RGFP966-treated rats, there was no significant change in PW1 amplitude in ABRs evoked by the extinguished 5.0 kHz signal tone or the non-extinguished 5.9 and 11.5 kHz tones (left panels). Further, among both (**A**) vehicle- and **(B)** RGFP966-treated rats, there was no difference in the amplitude shifts between pairs of frequencies (right panels). Bars indicate average change. Dots indicate individual data points.

**Figure 5.**
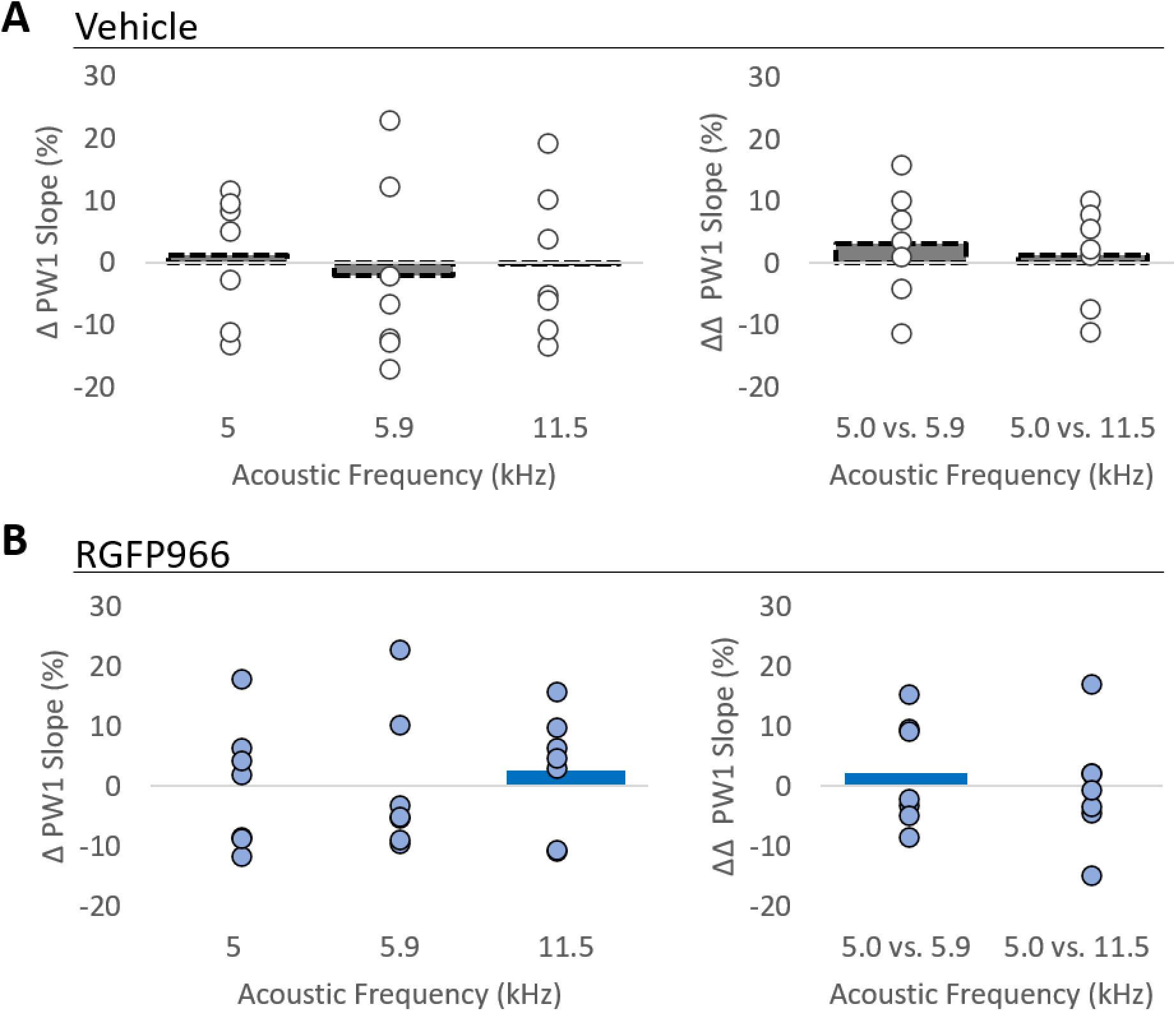
Single-tone extinction training does not result in PW1 slope changes in drug treatment groups. Among both (**A**) vehicle- and **(B)** RGFP966-treated rats, there was no significant change in PW1 slope in ABRs evoked by the extinguished 5.0 kHz signal tone or the non-extinguished 5.9 and 11.5 kHz tones (left panels). Further, among both (**A**) vehicle- and **(B)** RGFP966-treated rats, there was no difference in the slope shifts between pairs of frequencies (right panels). Bars indicate average change. Dots indicate individual data points.

**Figure 6.**
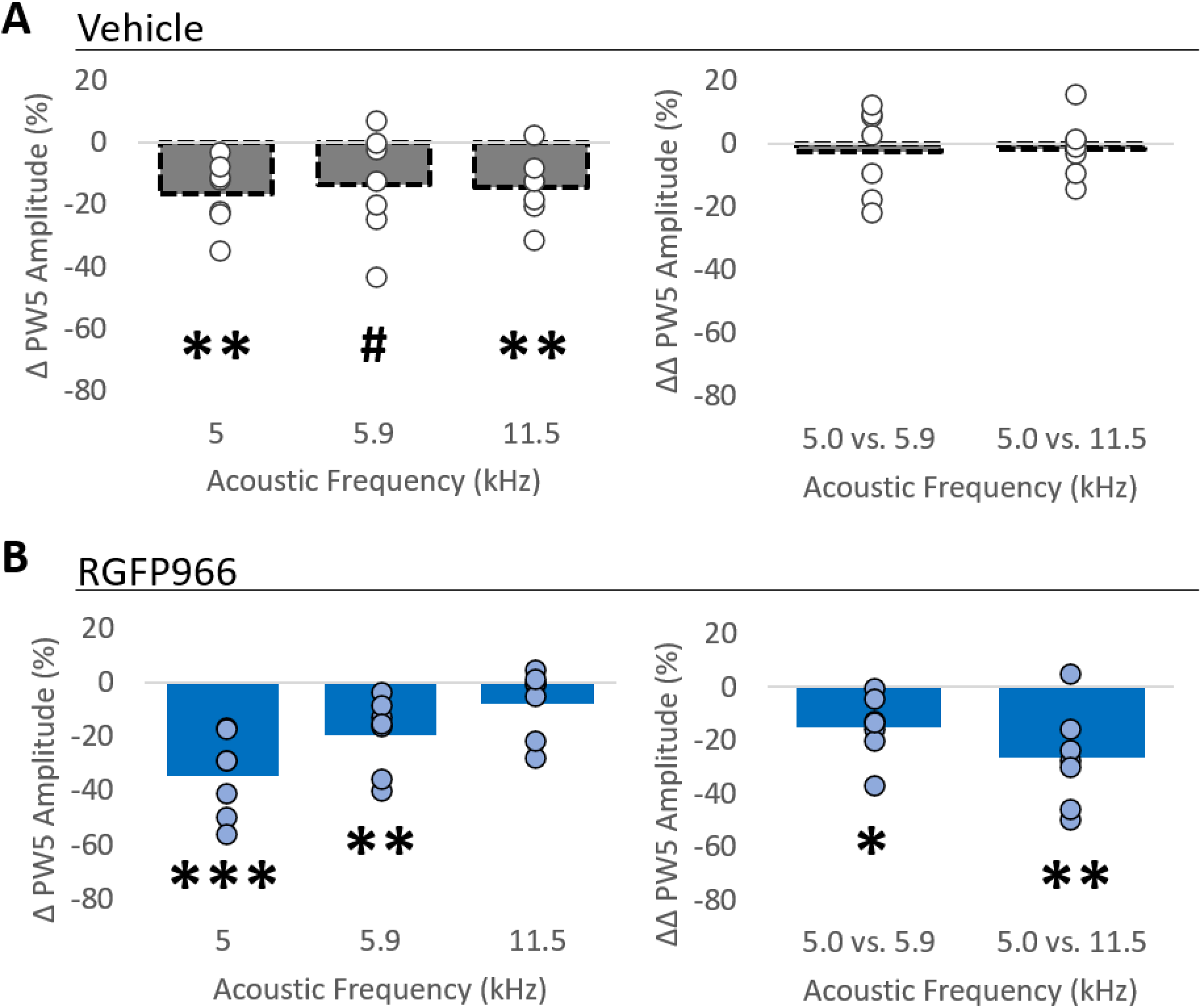
RGFP966-treatement is associated with larger decrease in PW5 amplitude from pre-to-post single-tone extinction training in ABRs evoked by the extinguished 5.0 kHz tone than in ABRs evoked by the non-extinguished tones. **(A)** Among vehicle-treated animals, there was a significant decrease from pre-to-post extinction training in PW5 amplitude in ABRs evoked by the 5.0 kHz and 11.5 kHz tones, and a trend toward a significant decrease in amplitude in ABRs evoked by the 5.946 kHz tone (left). There were no significant differences in the magnitude of PW5 amplitude change between pairs of frequencies (right). **(B)** Among RGFP966-treated animals, there was a significant decrease from pre-to-post extinction training in PW5 amplitude in ABRs evoked by the 5.0 and 5.946 kHz tones (left). However, the amplitude decrease in ABRs evoked by the 5.0 kHz extinguished tone was significantly *larger* than those in ABRs evoked by the non-extinguished 5.9 kHz tone and the non-extinguished 11.5 kHz tone (right). Bars indicate average change. Dots indicate individual data points. *p<0.05, **p<0.01, ***p<0.001, ^#^p<0.10

**Figure 7.**
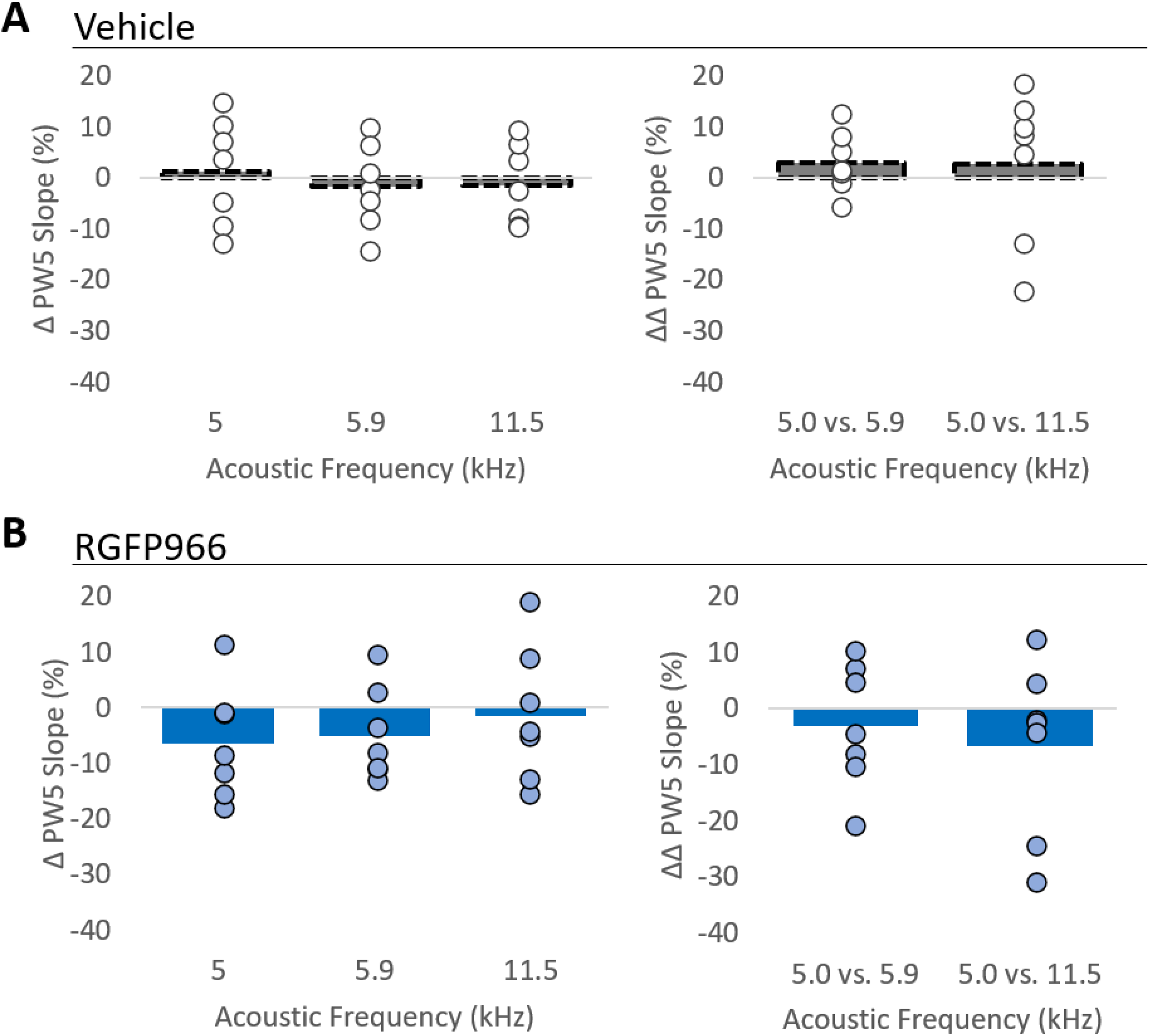
Single-tone extinction training does not result in PW5 slope changes in drug treatment groups. Among both (**A**) vehicle-treated and **(B)** RGFP966-treated rats, there were no significant changes in PW5 slope in ABRs evoked by the extinguished 5.0 kHz signal tone or the non-extinguished 5.9 and 11.5 kHz tones (left panels). Further, among both (**A**) vehicle-treated and **(B)** RGFP966-treated rats, there was no difference in the slope shift between pairs of frequencies (right panels). Bars indicate average change. Dots indicate individual data points.

**Figure 8.**
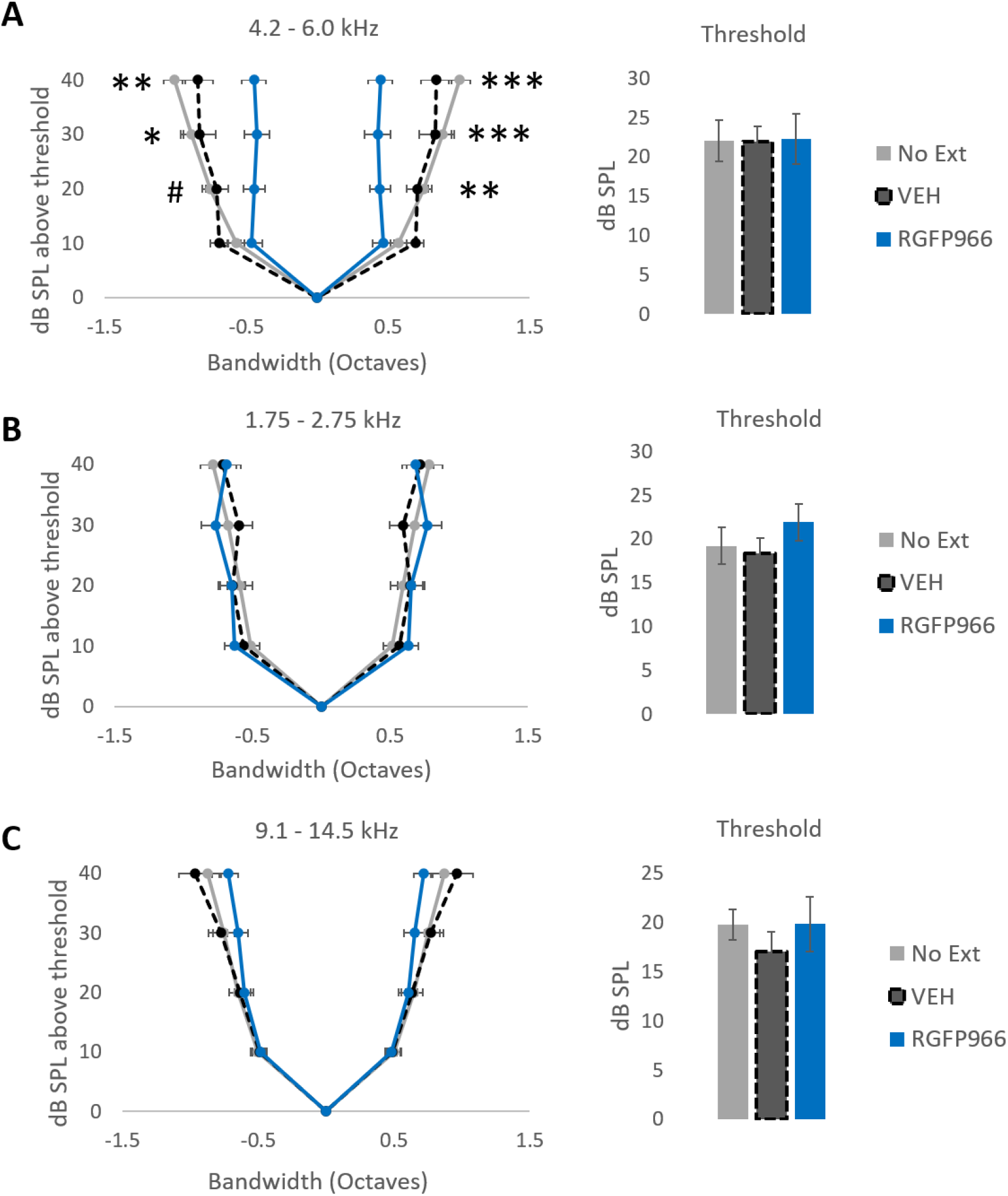
HDAC3 inhibition promotes learning-induced auditory cortical plasticity that is signal-specific. Panels represent tuning bandwidth and response threshold derived from sound-evoked neural responses from the auditory cortical recordings (**A**) Among auditory cortical sites tuned near the extinguished signal tone frequency (5.0 kHz; +/− 1/3 octave), RGFP966-treated animals (n= 5 animals, out of 7; 23 recording sites) showed narrower tuning bandwidth at multiple levels than vehicle-treated (n= 6 animals, out of 7; 27 recordings sites) and animals that did not receive extinction training (n = 6 animals; 37 recording sites). There were no group differences in response threshold (dB SPL). **(B)** Among auditory cortical sites tuned near the one of the non-extinguished signal tone frequencies (2.17 kHz; +/− 1/3 octave), there were no differences in tuning bandwidth between RGFP966-treated animals (n= 5 animals, out of 7; 27 recording sites), vehicle-treated animals (n=6 animals, out of 7; 22 recordings sites), and animals that did not receive extinction training (n=6 animals; 30 recording sites). There were no group differences in response threshold (dB SPL). **(C)** Among auditory cortical sites tuned near the one of the non-extinguished signal tone frequencies (11.5 kHz; +/− 1/3 octave), there were no differences in tuning bandwidth between RGFP966-treated animals (n= 5 animals, out of 7; 34 recording sites), vehicle-treated animals (n=6 animals, out of 7; 37 recordings sites), and animals that did not receive extinction training (n=6 animals; 57 recording sites). There were no group differences in response threshold (dB SPL). ***p<0.001, **p<0.01, *p<0.05, #p<0.10

**Figure 9.**
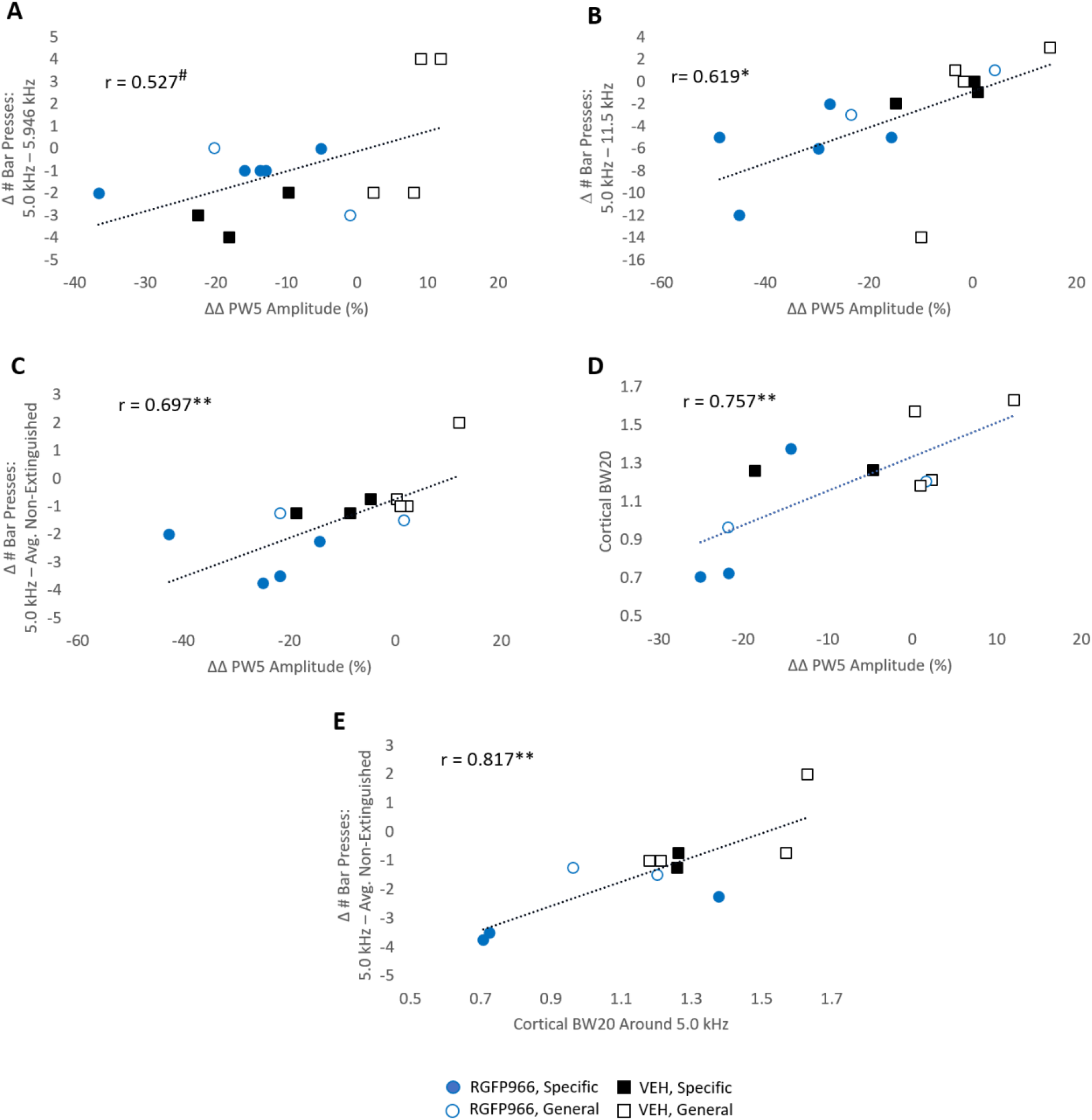
Auditory system plasticity is correlated with frequency-specific extinction memory. Each panel presents correlative data between two of three different types of measures: relative changes in PW5 amplitude (among pairs of frequencies; ΔΔPW5 Amplitude), behavioral contrast measures (among pairs of frequencies; Δ # Bar Presses), and auditory cortical tuning bandwidth (Cortical BW20) for sites tuned near the extinguished 5.0 kHz signal tone. **(A, B, C)** A greater PW5 amplitude decrease in ABRs evoked by 5.0 kHz (relative to non-extinguished tone frequencies) is associated with greater extinction memory specificity revealed behaviorally. **(D)** Greater PW5 amplitude decreases in ABRs evoked by 5.0 kHz (relative to non-extinguished tone frequencies) are associated with narrower auditory cortical tuning bandwidths (BW20) for sites tuned near the extinguished 5.0 kHz signal tone. **(E)** Narrower auditory cortical tuning bandwidth (BW20) for sites tuned near the extinguished 5.0 kHz signal tone is associated with greater extinction memory specificity. *p<0.05, **p<0.01, ^#^p<0.10

**Table 6.** Changes in PW1 following single-tone extinction training. This table presents change in PW1 slope and amplitude (M ± SE) from pre- to post-single-tone extinction training. Sample sizes are in parentheses.

|  |  | all | vehicle | RGFP966 | general | specific |
| --- | --- | --- | --- | --- | --- | --- |
| 5.0 kHz | amplitude | $-2.14 \pm 2.31$<br>(14) | $-3.09 \pm 2.95$<br>(7) | $-3.11 \pm 3.89$<br>(7) | $-1.05 \pm 3.09$<br>(6) | $-4.65 \pm 3.46$<br>(8) |
| | slope | $0.69 \pm 2.87$<br>(14) | $1.22 \pm 3.75$<br>(7) | $0.35 \pm 3.91$<br>(7) | $2.42 \pm 4.22$<br>(6) | $-0.44 \pm 3.47$<br>(8) |
| 5.946 kHz | amplitude | $-1.21 \pm 2.30$<br>(14) | $-1.33 \pm 3.42$<br>(7) | $-1.94 \pm 2.96$<br>(7) | $-0.73 \pm 3.97$<br>(6) | $-2.32 \pm 2.60$<br>(8) |
| | slope | $-1.74 \pm 3.52$<br>(14) | $-2.03 \pm 5.42$<br>(7) | $0.24 \pm 4.42$<br>(7) | $0.04 \pm 5.91$<br>(6) | $-1.6 \pm 4.26$<br>(8) |
| 11.5 kHz | amplitude | $-1.24 \pm 2.45$<br>(14) | $-2.49 \pm 3.03$<br>(7) | $-0.83 \pm 3.69$<br>(7) | $-0.31 \pm 3.42$<br>(6) | $-2.67 \pm 3.26$<br>(8) |
| | slope | $0.88 \pm 2.99$<br>(14) | $-0.13 \pm 4.39$<br>(7) | $2.67 \pm 3.70$<br>(7) | $4.46 \pm 4.44$<br>(6) | $-1.12 \pm 3.58$<br>(8) |

**Table 7.** Relative changes in PW1 between pairs of frequencies following single-tone extinction training. This table presents the relative difference in change in PW1 slope and amplitude (M ± SE) between pairs of frequencies from pre- to post-single-tone extinction training. Sample sizes are in parentheses.

|  |  | all | vehicle | RGFP966 | general | specific |
| --- | --- | --- | --- | --- | --- | --- |
| 5.0 vs.<br>5.946<br>kHz | amplitude | $-0.93 \pm 1.27$<br>(14) | $-1.76 \pm 1.61$<br>(7) | $-1.17 \pm 2.15$<br>(7) | $-0.31 \pm 1.86$<br>(6) | $-2.33 \pm 1.82$<br>(8) |
| | slope | $2.44 \pm 2.07$<br>(14) | $3.25 \pm 3.34$<br>(7) | $2.24 \pm 3.33$<br>(7) | $2.37 \pm 3.87$<br>(6) | $1.16 \pm 2.40$<br>(8) |
| 5.0 vs.<br>11.5 kHz | amplitude | $-0.89 \pm 1.84$<br>(14) | $-0.61 \pm 3.37$<br>(7) | $-2.29 \pm 1.52$<br>(7) | $-0.74 \pm 2.96$<br>(6) | $-1.98 \pm 2.36$<br>(8) |
| | slope | $-0.19 \pm 1.96$<br>(14) | $1.35 \pm 2.92$<br>(7) | $-0.19 \pm 3.54$<br>(7) | $-0.52 \pm 2.08$<br>(6) | $0.68 \pm 2.68$<br>(8) |

**Table 8.**
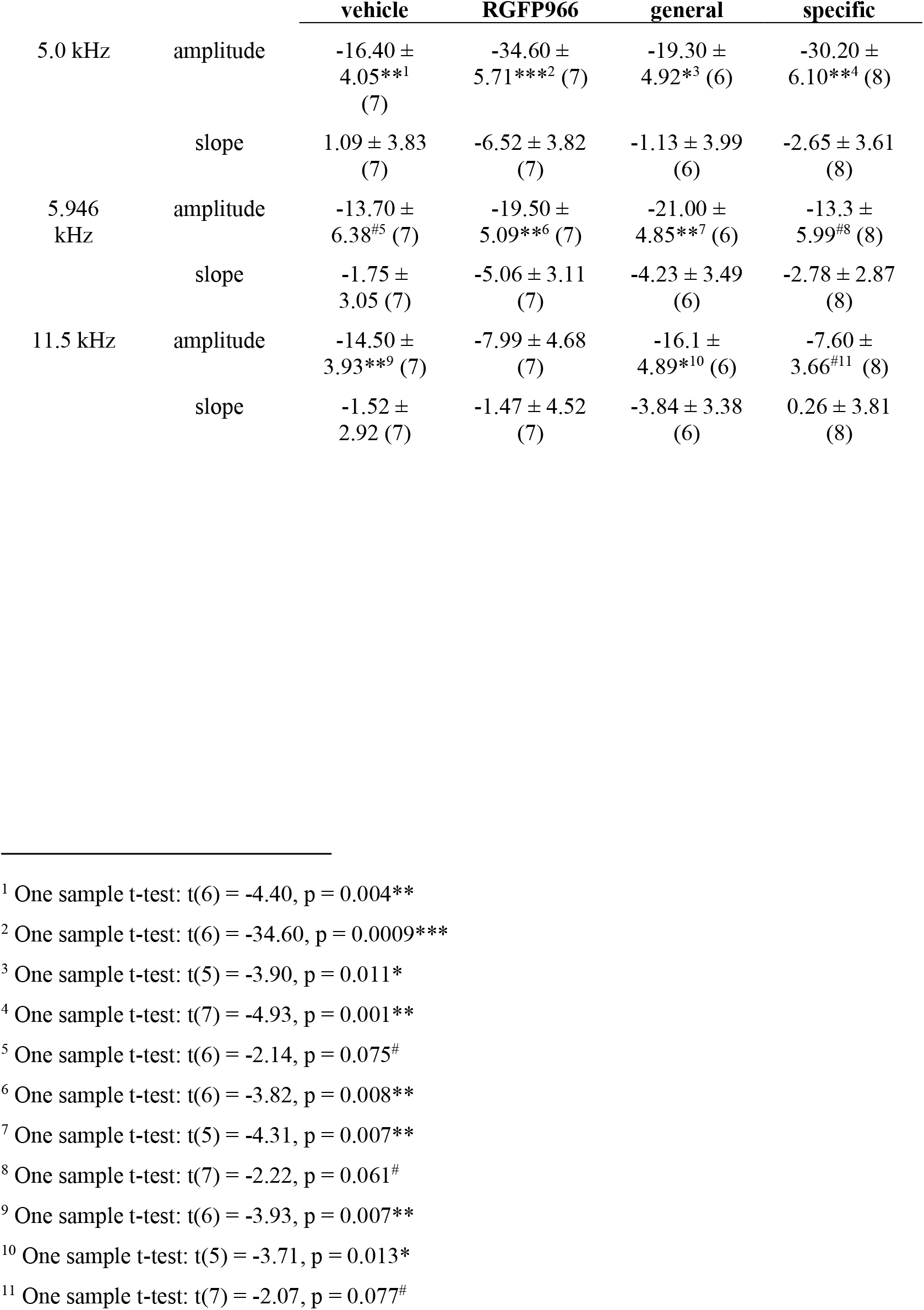
Changes in PW5 following single-tone extinction training. This table presents change in PW5 slope and amplitude (M ± SE) from pre- to post-single-tone extinction training. Sample sizes are in parentheses. *p<0.05, **p<0.01,***p<0.001, ^#^p<0.10

**Table 9.** Relative changes in PW5 between pairs of frequencies following single-tone extinction training. This table presents the relative difference in change in PW5 slope and amplitude (M ± SE) between pairs of frequencies from pre- to post-single-tone extinction training. Sample sizes are in parentheses. *p<0.05, **p<0.01, ^#^p<0.10

|  |  | vehicle | RGFP966 | general | specific |
| --- | --- | --- | --- | --- | --- |
| 5.0 vs.<br>5.946<br>kHz | amplitude | -2.71 ±<br>5.27 (7) | -15.10 ±<br>4.35*<sup>1</sup> (7) | 1.684 ±<br>4.77 (6) | -16.80 ±<br>3.38**<sup>2</sup> (8) |
| | slope | 2.843 ±<br>2.21 (7) | -3.19 ± 4.11<br>(7) | 3.09 ± 4.88<br>(6) | 0.137 ±<br>2.49 (8) |
| 5.0 vs.<br>11.5 kHz | amplitude | -1.92 ±<br>3.55 (7) | -26.60 ±<br>6.77**<sup>3</sup> (7) | -3.21 ± 5.29<br>(6) | -22.60 ±<br>6.63*<sup>4</sup> (8) |
| | slope | 2.614 ±<br>5.40 (7) | -6.78 ± 5.72<br>(7) | 2.709 ±<br>6.63 (6) | -2.91 ± 5.72<br>(8) |

<sup>1</sup> Holm-Bonferroni corrected paired samples t-test: t(6) = -3.465, p = 0.026\*
<sup>2</sup> Holm-Bonferroni corrected paired samples t-test: t(7) = -4.981, p = 0.001\*\*
<sup>3</sup> Holm-Bonferroni corrected paired samples t-test: t(6) = -3.930, p = 0.007\*\*
<sup>4</sup> Holm-Bonferroni corrected paired samples t-test: t(7) = -3.401, p = 0.022\*

### 3.4 HDAC3 Inhibition Promotes Frequency-Specific Auditory Cortical Plasticity

Frequency-specific memory for associations between tones and rewards, including memory enabled by HDAC3 inhibition, has been associated with signal-specific plasticity at the level of the primary auditory cortex (A1) (Recanzone, Schreiner, & Merzenich, 1993; Polley et al., 2006; Keeling et al., 2008; Bieszczad et al., 2010; Bieszczad & Weinberger, 2012; Bieszczad et al., 2015; Shang et al., 2019; Rotondo & Bieszczad, 2020). Previous studies reported that frequency-specific tone-reward memory is associated with a signal-specific sharpening of A1 tuning bandwidth (Bieszczad et al., 2015; Shang et al., 2019; Rotondo & Bieszczad, 2020). Here, electrophysiological recordings in A1 were following the Extinction Memory Test to compare forms of A1 plasticity in the treated groups (RGFP966 vs. vehicle). Electrophysiological data were analyzed according to the characteristic frequency of each site to parallel behavioral analyses by pooling neural data near the extinguished signal tone frequency (Fig. 8a) separately from sites tuned near the non-extinguished signal tone frequencies (Fig. 8b,c). Cortical receptive fields were analyzed for tuning bandwidth (BW, with respect to the breadth of frequency responsiveness in octaves) and response threshold (with respect to sound-evoked level in decibels). In sites tuned near the signal tone frequency (within one-third of an octave), RGFP966-treated animals had significantly narrower tuning bandwidth than vehicle-treated animals and animals that did not receive extinction training. (Table 10). In contrast, there were no differences in tuning bandwidth for sites tuned near (within one third of an octave) of the non-extinguished 2.17 kHz signal tone or the non-extinguished 11.5 kHz signal tone (Table 10). Cortical sound-evoked response threshold did not differ among groups (**sites tuned near 5.0 kHz**: RGFP966: n=23, M=22.25, SE=3.187; VEH: n=27, M=21.894, SE=2.028; No Ext: n=37, M=22.061, SE=2.580; one-way ANOVA: F(2,90) = 0.004, p=0.996; **sites tuned near 2.17 kHz:** RGFP966: n=28, M=21.93, SE=2.119 ; VEH: n=22, M=18.335, SE=1.8206; No Ext: n=30; M=19.241, SE=2.066; one-way ANOVA: F(2,79) = 0.813, p=0.447; **sites tuned near 11.5 kHz:** RGFP966: n=35, M=19.86, SE=2.775; VEH: n=40, M=17.082, SE=1.994; No Ext: n=37, M=19.832, SE=1.549; one-way ANOVA: F(2,131) = 0.595, p=0.553).

**Table 10.**
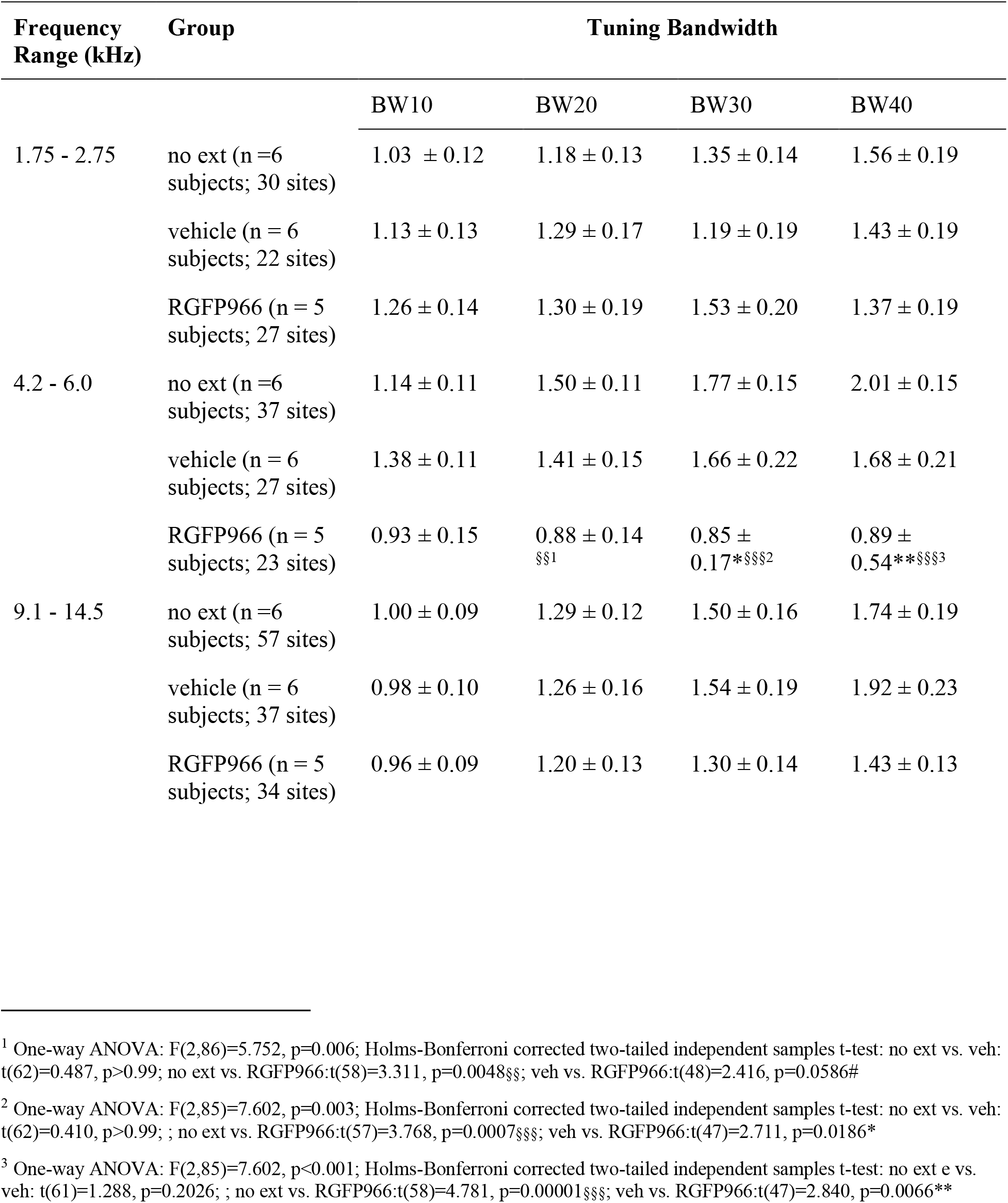
**C**ortical tuning bandwidth for RGFP966- and vehicle-treated animals. This table displays tuning bandwidth in octaves (M ± SE) for sites tuned around the 5.0 kHz extinguished tone (4.2-6.0 kHz), sites tuned around the non-extinguished 2.17 kHz tone (1.7-2.75), and sites tuned around the non-extinguished 11.5 kHz tone (9.1-14.5 kHz). *indicates a difference vs. vehicle-treated animals, where *p<0.05, **p<0.01, and ***p<0.001. § indicates a difference vs. no ext animals, §p<0.05, §§p<0.01, and §§§p<0.001.

Notably, re-grouping the cortical data by “specific” or “general” extinction memory yielded the similar pattern of results (Table 11). Animals with frequency-specific extinction memory exhibited narrower cortical tuning bandwidth around the extinguished 5.0 kHz signal tone than animals with frequency general memory and animals that did not receive extinction training. There were no group differences in tuning bandwidth around the non-extinguished 2.17 kHz tone or the non-extinguished 11.5 kHz tone (Table 11). There were no differences in response threshold (**sites tuned near 5.0 kHz**: specific: n=23, M=20.842, SE=2.869; general: n=27, M=23.66, SE=2.441; No Ext: n=37, M=22.061, SE=2.580; one-way ANOVA: F(2,87) = 0.247, p=0.782; **sites tuned near 2.17 kHz**: specific: n=27, M=19.888, SE=2.146; general: n=23, M=20.890, SE=1.906;No Ext: n=30; M=19.241, SE=2.066; one-way ANOVA: F(2,79) = 0.156, p=0.856; **sites tuned near 11.5 kHz**: specific: n=36, M=17.258, SE=2.575; general: n=35, M=19.74, SE=2.270; No Ext: n=37, M=19.832, SE=1.549; one-way ANOVA: F(2,131) = 0.472, p=0.625).

**Table 11.**
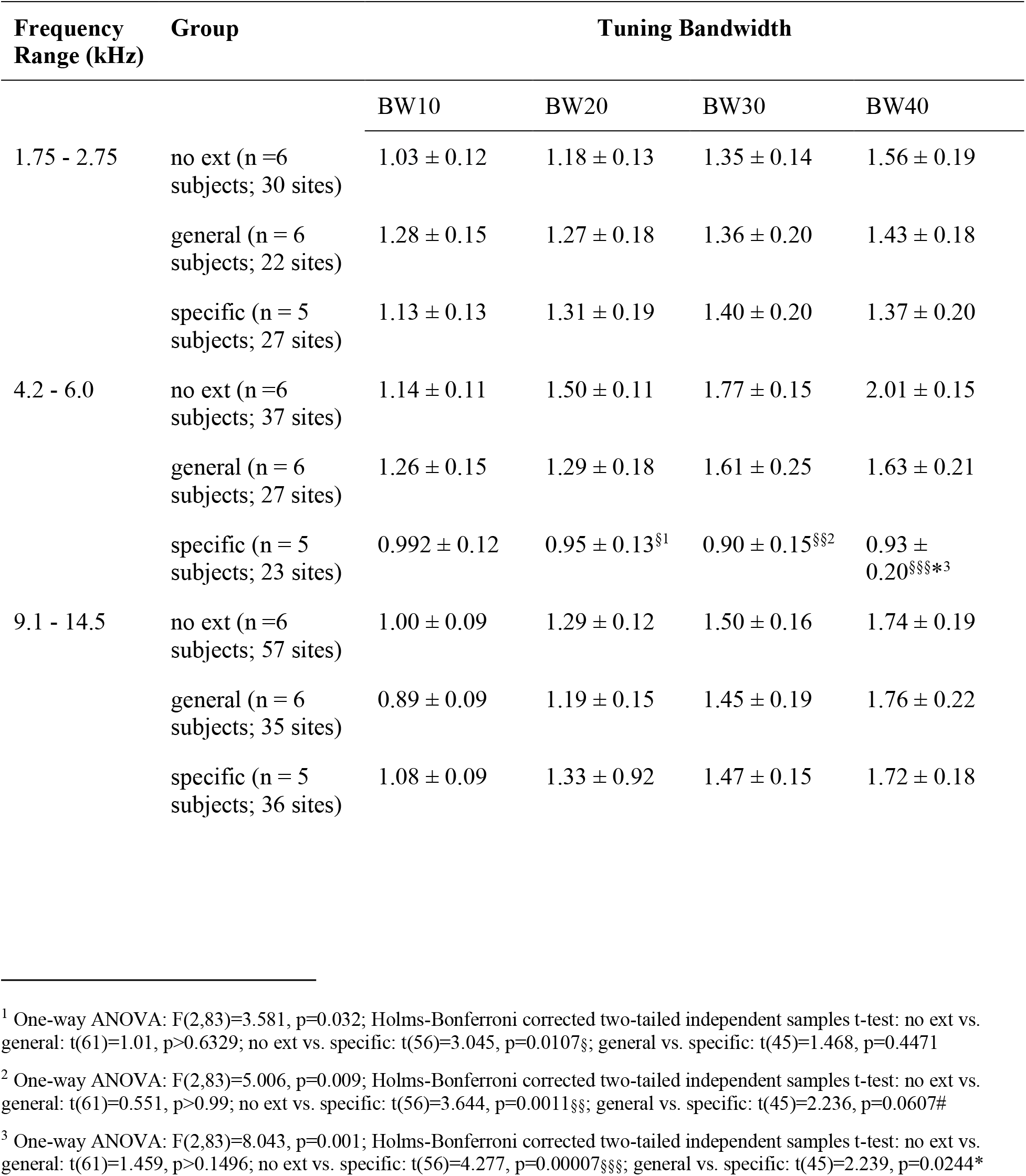
Cortical tuning bandwidth for animals with frequency-specific and -general extinction memories. This table displays tuning bandwidth (M ± SE) for sites tuned around the 5.0 kHz extinguished tone (4.2-6.0 kHz), sites tuned around the non-extinguished 2.17 kHz tone (1.7-2.75), and sites tuned around the non-extinguished 11.5 kHz tone (9.1-14.5 kHz). *indicates a difference vs. animals with general memories, where *p<0.05, **p<0.01, and ***p<0.001. § indicates a difference vs. no ext animals, §p<0.05, §§p<0.01, and §§§p<0.001.

### 3.5 Auditory System Plasticity is Correlated with Individual Differences in Extinction Memory

To determine whether changes in sound-evoked auditory processing were meaningfully related to individual differences in learned behavior, measures of neurophysiological plasticity that showed significant learning-induced changes were correlated with behavioral at the time of extinction training and the extinction memory test.

Despite observing significant increases in PW5 amplitude and slope due to multi-tone reward training, these changes did not correlate with behavior at the time of single-tone extinction training, including the number of bar-presses to 5.0 kHz during the first extinction session (amplitude: n = 13, r = 0.076, p – 0.805; slope: n = 13, r = 0.417, p=0.156) or bar-presses to 5.0 kHz during the second extinction session, expressed as a proportion of bar-presses during session 1 (amplitude: n = 13, r = 0.220, p = 0.470; slope: n = 13, r = −0.114, p = 0.718).

In contrast, a number of significant relationships were found between measures of extinction-induced auditory system plasticity and extinction memory specificity (Fig. 9; Table 12). Memory specificity here is defined with respect to relative differences in behavioral responding to the extinguished tone vs. at least one other non-extinguished tone (as described in *2.6.1 Behavioral Statistical Analysis*). There was a significant positive relationship in the behavioral contrast between the extinguished 5.0 kHz tone and the non-extinguished 11.5 kHz tone (# bar presses to 5.0 kHz - # bar presses to 11.5 kHz) at the Extinction Memory Test and the relative difference in the change (between 5.0 kHz and 11.5 kHz) of PW5 amplitude: Animals that made relatively *fewer* responses to the extinguished 5.0 kHz tone also had a relatively *larger decrease* in PW5 amplitude in ABRs evoked by the 5.0 kHz tone (Fig. 9b). Similarly, there was a trend toward a significant positive relationship in the behavioral contrast between the extinguished 5.0 kHz tone and the non-extinguished 5.946 kHz tone (# bar presses to 5.0 kHz - # bar presses to 5.946 kHz) at the Extinction Memory Test and the relative difference in the change (between 5.0 kHz and 5.946 kHz) of PW5 amplitude: Animals that made relatively *fewer* response to the extinguished 5.0 kHz tone (vs. 5.946 kHz) also had a relatively *larger decrease* in PW5 amplitude in ABRs evoked by the 5.0 kHz tone (vs. 5.946 kHz) (Fig. 9a). Additionally, a “total” behavioral specificity measure (((# BPs to 5.0 kHz - # BPs to Nearby Tones) + (# BPs to 5.0 kHz - #BPs to Distant Tones)) / 2) had a significant positive relationship with the average relative difference in the change in PW5 amplitude of 5.0 vs. 5.946 kHz and 5.0 vs. 11.5 kHz: Animals that made relatively *fewer* responses to 5.0 kHz (vs. all other non-extinguished tones) also had a relatively greater average decrease in PW5 amplitude in ABRs evoked by the 5.0 kHz tone (vs. other tones) (Fig. 9c).

**Table 12.** Summary of Correlations with Extinction-Induced Changes in PW5 Amplitude. This table displays the Pearson r values for correlative data between extinction induced changes in PW5 amplitude (in ABRs evoked by the extinguished 5.0 kHz tone vs. non-extinguished tones), behavioral contrast scores among pairs of frequencies during the Extinction Memory Test, and auditory cortical bandwidth (BW20) for sites tuned near the extinguished tone frequency. *p<0.05, **p<0.01, ***p<0.001, ^#^p<0.10; ΔΔ: difference in the post-pre extinction change; Δ%: difference in percentage

|  | **All** | **Vehicle** | **RGFP966** | **General** | **Specific** |
| --- | --- | --- | --- | --- | --- |
| 5.0 - 5.9 kHz:<br>ΔΔ ABR vs.<br>Δ% Bar presses | 0.527<sup>#1</sup><br>(14) | 0.751<sup>#2</sup><br>(7) | 0.024<br>(7) | 0.350<br>(6) | 0.459<br>(8) |
| 5.0 – 11.5 kHz:<br>ΔΔ ABR vs.<br>Δ% Bar presses | 0.619<sup>*3</sup><br>(14) | 0.584<br>(7) | 0.739<sup>#4</sup><br>(7) | 0.538<br>(6) | 0.758<sup>*5</sup><br>(8) |
| 5.0 – (avg. of 5.9 &<br>11.5 kHz):<br>ΔΔ ABR vs.<br>Δ% Bar presses | 0.697<sup>**6</sup><br>(14) | 0.583<br>(7) | 0.483<br>(7) | 0.590<br>(6) | 0.441<br>(8) |
| BW20 vs. ΔΔ ABR<br>(5.0 – (avg. of 5.9<br>& 11.5 kHz)) | 0.757<sup>**7</sup><br>(11) | 0.519<br>(6) | 0.684<br>(5) | 0.763<sup>#8</sup><br>(6) | 0.728<br>(5) |
| BW20 vs. Δ% Bar<br>presses (5.0 vs. avg.<br>of non-extinguished<br>tones) | 0.817<sup>**9</sup><br>(11) | 0.738<sup>#10</sup><br>(6) | 0.590<br>(5) | 0.602<br>(6) | 0.736<br>(5) |

<sup>1</sup> p = 0.052
<sup>2</sup> p = 0.051
<sup>3</sup> p = 0.018
<sup>4</sup> p = 0.057
<sup>5</sup> p = 0.029
<sup>6</sup> p = 0.005
<sup>7</sup> p = 0.006
<sup>8</sup> p = 0.077
<sup>9</sup> p = 0.002

Notably, changes in PW5 due to multi-tone reward training did not have any predictive value for subsequent behavior during the Extinction Memory Test. Specifically, the behavioral contrast between the extinguished 5.0 kHz tone and the non-extinguished 5.946 kHz tone (# bar presses to 5.0 kHz - # bar presses to 5.946 kHz) was not related to the relative difference in the change (between 5.0 kHz and 5.946 kHz) of PW5 amplitude (n =13, r = 0.106, p = 0.730) or PW5 slope (n = 13, r = 0.401, p = 0.174). Similarly, the behavioral contrast between the extinguished 5.0 kHz tone and the non-extinguished 11.5 kHz tone (# bar presses to 5.0 kHz - # bar presses to 11.5 kHz) was not related to the relative difference in the change (between 5.0 kHz and 11.5 kHz) of PW5 amplitude (n =13, r = 0.388, p = 0.170) or PW5 slope (n = 13, r = 0.100, p = 0.733). Moreover, the relative changes in PW5 (5.0 vs 5.946 kHz and 5.0 vs 11.5 kHz) due to multi-tone reward training did not correlate with cortical tuning bandwidth measured after extinction training (amplitude: n= 10, r = 0.178, p = 0.622; slope: n = 10, r = 0.223, p = 0.535). Therefore, extinction driven, and not tone-reward, effects on subcortical plasticity are predictive of extinction memory specificity.

Therefore, we investigated the relationship between cortical and subcortical plasticity induced by extinction. Finally, there was a significant positive correlation between the average relative difference in the change in PW5 amplitude of 5.0 vs. 5.946 kHz and 5.0 vs. 11.5 kHz and auditory cortical bandwidth for sites tuned near 5.0 kHz: Individuals that had relatively *larger decreases* in PW5 amplitude in ABRs evoked by the 5.0 kHz tone also exhibited *narrower cortical tuning bandwidth* in A1 sites tuned near the 5.0 kHz frequency. Finally, these data validate the relationship between frequency-specific memory and narrowed cortical bandwidth tuning previously reported for initial tone-reward memory (Rotondo & Bieszczad, 2020). There was a significant correlation between cortical bandwidth for sites tuned near the extinguished signal tone and extinction memory specificity: Individuals with more specific extinction memory exhibited narrower cortical tuning bandwidth for the signal tone. In sum, these findings support that the link between HDAC3 and memory specificity may be mediated by signal-specific auditory neuroplasticity in the auditory cortical and subcortical systems.

## 4 Discussion

We report the first investigations of cue-specific auditory extinction memory, the effects of HDAC3 inhibition (HDAC3-i) on extinction memory specificity, and the auditory neural substrates of frequency-specific extinction memory. Data support that HDAC3-i enhances the frequency-specificity of extinction memory. RGFP966 administered immediately after extinction sessions resulted in the inhibition of behavioral responding to the extinguished frequency signal more than to non-extinguished frequencies. Extinction memory specificity was characterized neurophysiologically by signal-specific changes in auditory cortical plasticity in the form of sharpened frequency tuning bandwidth, and for the first time, signal-specific auditory subcortical plasticity in the form of amplitude decreases in PW5 of the ABR. Significant brain-behavior relationships validated that the detected forms of signal-specific auditory plasticity are a neural substrate of frequency-specific extinction memory and behavior. Taken together with previous studies of the effects of HDAC3 inhibition on auditory memory specificity in reward learning and discrimination learning (Rotondo & Bieszczad, 2020; Shang et al., 2019; Bieszczad et al. 2015), the results suggest that: (1) HDAC3-i acts on mechanisms that encode specific sensory details into memory *regardless of associative value and behavioral value*, and (2) sensory signal-specificity *per se* is encoded in memory by signal-specific changes in sensory system processing of behaviorally-relevant sound features.

Epigenetic mechanisms like HDACs have been implicated in memory in a variety of tasks (Vecsey et al., 2007; Stefanko et al., 2009; McQuown et al., 2011; Malvaez et al., 2013; Phan et al., 2017). Present results support this fundamental role for HDAC3-mediated regulation of memory consolidation. Previous literature has focused on the ability of HDAC3 inhibition to enhance the formation, strength, or persistence of long-term memory (Vecsey et al., 2007; Stefanko et al., 2009), including extinction memory (Wang et al., 2010; Malvaez et al., 2013; Castino et al., 2015; Hitchcock et al., 2019). Yet, some studies failed to find effects on memory strength without consideration of memory specificity first (Bieszczad et al., 2015, Shang et al., 2019, Rotondo & Bieszczad, 2020), including in auditory fear extinction memory (Bredy et al., 2007; Bowers et al, 2015). However, those that have *also* probed the sensory specificity of memory have reported that HDAC3-i can enhance specificity even in absence of effects on other cardinal features of memory like its strength or persistence. Thus, future studies should consider that memory has many characteristic features and determine exactly which features and their underlying mechanisms are altered by epigenetic manipulations. The present study represents an important step in revealing that HDAC3-i enhances sensory signal-specificity regardless of the phase of learning and the direction of learned behavior.

Importantly, the effect of HDAC3-i on auditory system plasticity mimics the pattern of naturally-formed frequency-specific extinction memory. Grouping neurophysiological data by treatment group *or* by extinction memory phenotype (frequency-general vs. –specific) produces the same pattern of results: frequency-specific extinction memory is associated with signal-specific auditory system plasticity. Thus, not only does the present study reveal that HDAC3 is a mechanism of gating signal-specific plasticity, and subsequently, signal-specific extinction memory; it provides information on the nature of plasticity as a neural coding mechanism that supports sensory specificity in memory.

### Frequency-specific memory is associated with signal specific plasticity regardless of task type

Frequency-specific extinction memory associated with signal-specific auditory cortical and subcortical plasticity replicates the pattern of plasticity associated with frequency-specific memory in a tone-reward training task (Bieszczad et al., 2015; Rotondo & Bieszczad, 2020). Signal-specific narrowing of auditory cortical tuning bandwidth is in agreement with studies demonstrating that plasticity in auditory cortical frequency representations reflects what is actually learned about sound features, rather than pure stimulus statistics of the acoustic experience (Polley et al., 2006; Weinberger et al., 2013). Further, changes in the ABR validate that signal-specific auditory subcortical plasticity is associated with frequency-specific memory. First, only individuals that form frequency-specific extinction memory develop signal-specific subcortical plasticity in the form of decreased PW5 amplitude. This is consistent with individuals who form frequency-specific tone-reward memory after an initial phase of learning (Rotondo & Bieszczad, 2020). Second, when subjects are trained to respond generally across frequency (i.e., in multi-tone reward training) signal-specific plasticity does not develop; it is either not detected (e.g., in PW1) or develops the same way across frequencies (e.g., in PW5). The selectivity of plasticity for behaviorally-salient signals has been proposed to stem from interaction between the auditory cortex and subcortex (Strait et al., 2012; Chandrasekaran et al., 2014), which is supported here by significant correlations in the magnitude of plasticity between these levels of the auditory system, as well as between auditory system plasticity and signal-specific behavior.

### Frequency-specific plasticity emerges both cortically and subcortically with frequency-specific behavior, regardless of the behavioral direction of response that is cued by a sound signal

Because extinction represents a reversal of sound meaning and behavior from ‘original’ sound-reward learning, one prediction is that the direction of plasticity observed in auditory extinction will reverse that of tone-reward learning. Previous studies have supported this in the auditory system (Disterhoft & Stuart, 1976; Lai, Adler, & Gan, 2018), though sometimes the reversal continues beyond the point of the naïve state to imply an opposing rather than a purely reversing process (Bieszczad & Weinberger, 2012). However, in other regions, extinction learning entails the induction of new, distinct forms of plasticity that might be necessary to update links between cues to newly appropriate behaviors (Corcoran et al., 2005; Sepulveda-Orenga et al., 2013). Here, we argue that forms of sensory system plasticity that primarily function to code for frequency-specificity will not reverse after extinction, whereas forms of plasticity that code for other cardinal features of memory will show a reversal.

Consistent with previous reports that extinction induces reversals of initial learning-induced gains (Disterhoft & Stuart, 1976), we report PW5 amplitude changes reverse with extinction in the form of an amplitude decrease. The *relative* amplitude decrease between ABRs evoked by extinguished vs. non-extinguished tones correlated with frequency-specific extinction memory: the greater the extinguished frequency-specific amplitude, the more specific the extinction to the extinguished frequency. Relative PW5 amplitude decreases after extinction training were actually *predictive* of differences in behavioral responding between pairs of frequencies during extinction memory testing. Therefore, changes in PW5 amplitude may contribute to coding frequency-specific extinction memory. Moreover, given that the direction of amplitude change increased in the initial training phase and *reversed* with extinction, this form of plasticity may additionally code for other aspects of memory, like the link from the cue to initiate (by an amplitude increase) vs. inhibit (by an amplitude decrease) behavioral responses.

In contrast to PW5 amplitude, PW5 slope plasticity induced by tone-reward training exhibited a resistance to further plasticity with extinction training. This may be interpreted in the context of extinction as new learning with its own (partially) unique set of neural changes. Alternatively, learning-induced plasticity that is resistant to change may be an important factor underlying behavioral recovery processes following extinction (Bouton, 2004). The lack of signal-specific plasticity of the PW5 slope in the present experiment does not rule out that it could be sensitive to learning-induced plasticity. In fact, the control analysis using animals that learned a single tone-reward association with frequency-specificity do show signal-specific changes in PW5 slope. Thus, one possible explanation is that signal-specific PW5 slope plasticity emerges with extended training and would have been observed in the present study with more extensive extinction training. However, the present data do not allow the conclusion that PW5 slope encodes specificity, regardless of task or phase of training, *per se*.

Importantly, frequency-specific extinction memory was also associated with auditory cortical plasticity that is likewise signal-specific. Moreover, frequency-specific sharpening of tuning bandwidth developed only after extinction learning. In contrast, subjects with frequency-general memory after extinction training show no change in tuning bandwidth, just as in subjects with frequency-general memory in an initial phase of sound-reward learning do not exhibit narrowed tuning (Rotondo & Bieszczad 2020). Frequency-general memory after single-tone extinction leaves tuning as broad as control subjects that did not receive extinction training at all. This is a particularly notable finding because the direction of cortical bandwidth plasticity (i.e., narrowing tuning) is the same as that observed with an initial frequency-specific tone-reward memory (Bieszczad et al.,2015; Shang et al., 2019; Rotondo & Bieszczad, 2020). The relationship between receptive field breadth and memory specificity has been observed even in other sensory systems (Kass et al., 2013; Kass & McGann, 2017). Thus, the present data support that narrowed sensory receptive field tuning may code for memory specificity *per se*, regardless of the type of associative learning task, or the ‘meaning’ of the sensory stimulus, or the learned behavioral output.

In sum, the present study shows that epigenetic manipulation, such as by HDAC3-i, is a valuable tool to understand sensory codes underlying features of memory. HDAC3-i may confer memory specificity at least in part by acting on naturally occurring auditory substrates that modify the representation of behaviorally relevant stimuli in a signal-specific manner. Learning-induced signal-specific sensory changes may contribute to the larger network of brain regions that support memory formation by specifying the identity of behaviorally relevant cues.

## ACKNOWLEDGEMENTS

This work was supported by NIH R03-DC014753 (to K.M.B.) and American Speech-Hearing-Language Foundation 2017 New Century Scholars Grant (to K.M.B.) and The Brain & Behavior Foundation 2017 NARSAD Young Investigator Award (to K.M.B.).

